# Bioinformatic prediction and high throughput in vivo screening to identify cis-regulatory elements for the development of algal synthetic promoters

**DOI:** 10.1101/2024.03.20.585996

**Authors:** Y. Torres-Tiji, H. Sethuram, A. Gupta, J. McCauley, J.V. Dutra-Molino, R. Pathania, L. Saxton, K. Kang, N. J. Hillson, S. P. Mayfield

## Abstract

Algae biotechnology holds immense promise for revolutionizing the bioeconomy through the sustainable and scalable production of various bioproducts. However, its development has been hindered by the lack of advanced genetic tools. This study introduces a synthetic biology approach to develop such tools, focusing on the construction and testing of synthetic promoters. By analyzing conserved DNA motifs within the promoter regions of highly expressed genes across six different algal species, we identified cis-regulatory elements (CREs) associated with high transcriptional activity. Combining the algorithms POWRS, STREME and PhyloGibbs, we predicted 1511 CREs and inserted them into a minimal synthetic promoter sequence in 1, 2 or 3 copies, resulting in 4533 distinct synthetic promoters. These promoters were evaluated in vivo for their capacity to drive the expression of a transgene in a high-throughput manner through next-generation sequencing post antibiotic selection and fluorescence-activated cell sorting. To validate our approach, we sequenced hundreds of transgenic lines showing high GFP expression. Further, we individually tested fourteen identified promoters, revealing substantial increases in GFP expression—up to nine times higher than the baseline synthetic promoter, with five matching or even surpassing the performance of the native AR1 promoter. As a result of this study, we identified a catalog of CREs that can now be used to build superior synthetic algal promoters. More importantly, here we present a validated pipeline to generate building blocks for innovative synthetic genetic tools applicable to any algal species with a sequenced genome and transcriptome dataset.

**Graphical Abstract:** 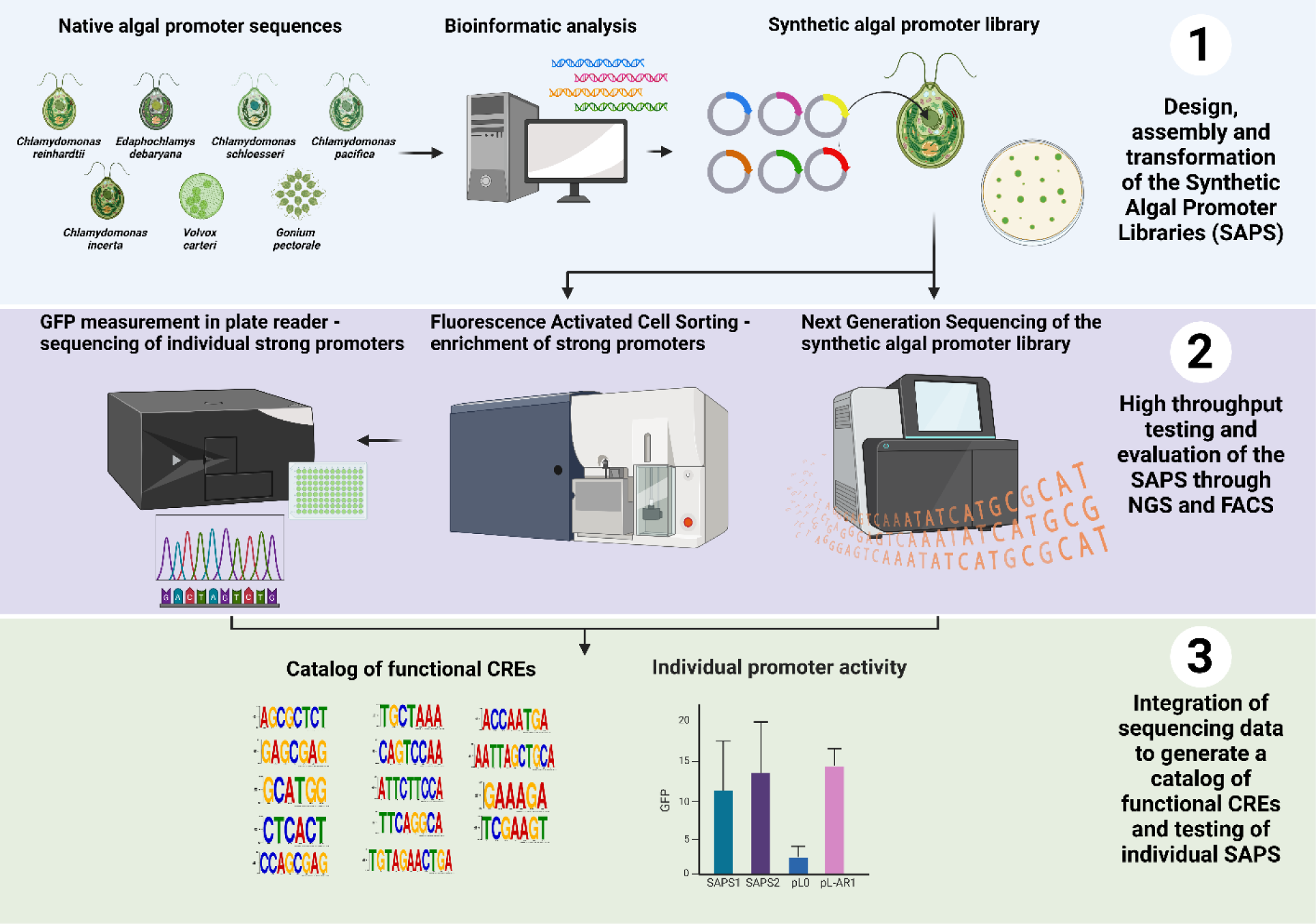

## Introduction

Biotechnology, the use of biological organisms and processes to create products, has ancient roots dating back 10,000 years with the advent of agriculture, and at least 6,000 years ago in fermenting beverages and making cheese^1^. However, the modern era of biotechnology began in the 1970s with the development of molecular cloning technologies, leading to a new industry, which was first exemplified by companies like Genentech, Amgen, and Chiron. Today, biotechnology significantly impacts the world’s economy, and is predicted to reach almost 14% of the global economy by the year 2030 ^2^.

As the biotechnology industry has matured, it has become clear that more sustainable and efficient organisms need to be developed, ones that can meet the ever-growing demand for new bio-based products, without overtaxing our already strained resources, like potable water and arable land ^3^. Microalgae, microscopic aquatic organisms, are poised to become a cornerstone of future sustainable biotechnology, as these versatile organisms can be engineered for the efficient and environmentally favorable production of food and feed, therapeutics, biofuels, bioplastics, and many other products ^3^. Numerous examples of current algae-based products exist, including omega-3 fatty acids and other nutritional supplements marketed for their health benefits, biopolymers, alternative food, and even biofuels ^4^.

However, optimizing the production of these valuable products requires precise control over the genetics and metabolism of these organisms, which can be achieved today through targeted genetic modifications, but at a much less sophisticated level than for other industrial biotechnology platforms. Fine-tuning metabolic pathways and expressing specific genes can significantly enhance the yield and quality of desired products, and to achieve this will require the development of sophisticated genetic tools, including promoters capable of driving robust and tunable transgene expression. Currently available genetic tools for algae fall short of those available in other model organisms like bacteria and yeast ^5^. While native promoters from the model green algae *Chlamydomonas reinhardtii*, such as the chimeric promoter AR1^6^, have been successfully utilized, their effectiveness pales compared to the diverse and powerful tools available in established industrial organisms.

Synthetic biology offers a promising avenue for overcoming these limitations and creating next-generation genetic tools for algae. This field of research focuses on designing and building biological systems with specific functions, and it has already yielded impressive results in other model organisms^7 8^. In the algae biotechnology field, synthetic biology is still in its initial stages. The first synthetic promoters in *C. reinhardtii* were created by analyzing the promoter regions of the 50 highest expressed genes in *C. reinhardtii* and identifying DNA motifs based on the presence of such elements in these genes’ promoters, and on their positional relationship within the promoter region, using the POWRS algorithm ^9^. Using this data, 25 synthetic algal promoters were made and the performance of which was compared to native hybrid promoter AR1 using in vivo expression assays, with seven of the synthetic promoters outperforming AR1, and one motif or CRE (CCCAT motif) was identified as a main driver of expression in one of the promoters ^10^. In a different study, some of the motifs identified by Scranton et al. were used to design better synthetic promoters by inserting them into native chimeric promoters ^11^. In another study, promoter regions of highly expressed genes were analyzed using software like WEEDER^12^, HOMER^13^, DREME^14^, and MEME^15^, followed by motif clustering and enrichment analysis. This process led to the identification of 13 putative DNA motifs that could function as transcriptional enhancers, which were synthesized and tested in vivo for their ability to drive the expression of a yellow fluorescent protein reporter gene. Out of these, five DNA motifs showed significantly higher reporter expression compared to AR1 ^16^. These studies have proven that it is possible to identify cis-regulatory motifs and utilize them to design functional synthetic promoters. However, the number of identified CREs remains limited, and the transcriptional activities of these elements have not been well characterized.

In this study, we sought to identify a large number of CREs by analyzing conserved DNA elements in the promoter regions of highly expressed genes across six different algal species. Using different bioinformatic algorithms, two pools of CREs were predicted, one containing 680 and the other 831 CREs. Consequently, two synthetic algal promoter (SAPS) libraries were built, with each CRE being tested by inserting 1, 2, or 3 tandem copies of a single CRE per synthetic promoter, thus generating 2040 promoters in SAPS1 and 2493 in SAPS2. We then transformed the promoter libraries into *C. reinhardtii* and assessed the impact of these CREs on gene expression using an in vivo reporter assay based on antibiotic selection and fluorescent protein gene expression, followed by Next-Generation Sequencing (NGS) to identify individual CREs. This enabled a high throughput comparison of relative promoter activity within a population containing thousands of transgenic lines to determine a relative ranking of these CREs. The sequencing data derived from the NGS analysis allowed us to create a catalog of more than 100 functional CREs in *C. reinhardtii* that can be used to generate functional synthetic algal promoter. Finally, we isolated 14 individual transformants containing synthetic promoters capable of driving Green Fluorescent Protein (GFP) at high levels through Fluorescence Activated Cell Sorting (FACS), and quantitated GFP expression from these promoters in detail, to demonstrate the utility of such synthetic promoters for driving recombinant gene expression.

## Results

### Bioinformatic analysis of conserved DNA motifs in the promoter regions of the highest expressed genes across different algal species

In this study, we identified a comprehensive list of CREs from algal genomes which can promote transgene expression upon insertion into synthetic promoters. Our hypothesis was that promoters exhibiting high gene expression possess DNA binding sites for transcription factors which, upon binding, augment transcriptional activity. We anticipated these DNA binding sites to be overrepresented in the promoter regions of genes with high expression levels. Furthermore, we expected these functional DNA sequences to be conserved across the promoter regions of orthologous genes in closely related species. Acknowledging the significant variability in motif prediction accuracy across datasets and species highlighted in prior studies, we elected a strategy that involved the application of three distinct motif prediction algorithms to identify CREs in *C. reinhardtii*^17^. Given these criteria, we employed the algorithms POWRS and STREME, in SAPS1, and PhyloGibbs, in SAPS2, to predict a total of 1471 potential CREs.

For SAPS1, the top 146 highly expressed genes in *C. reinhardtii* and their orthologues in *C. incerta* and *C. pacifica* were used as input sequences, anticipating an overrepresentation of transcriptionally activating CREs. Our constraints were kept lenient to minimize the number of false negatives, and we were confident in our high-throughput screening methods to manage even substantial quantities of false positives. To filter out ubiquitous DNA motifs in promoters unrelated to high transcriptional activity, control sequences were derived from the 146 least expressed genes in *C. reinhardtii* and *C. pacifica*. The input sequences are available in the Supporting Information – S1. The *POWRS* (POsition-sensitive WoRd Set) is a discriminative method that evaluates the significance of a motif through binomial distribution analysis of input and control sequences. Unlike other approaches, POWRS considers the motif’s position in the sequences, which is often overlooked, without resorting to Position Weight Matrices. Instead, it represents motifs using a single, non-degenerate consensus sequence along with some variants that differ by only one base pair. *STREME* (Simple, Thorough, Rapid, Enriched Motif Elicitation), a widely popular tool designed for ungapped motif discovery, uses Fisher’s Exact Test or the Binomial test to determine motif significance within the input sequences set relative to the control sequences. Only the motifs found in at least two of the three species were selected. This resulted in the prediction of 640 different CREs (Figure 1 and Supporting Information – S2).

**Figure 1:**
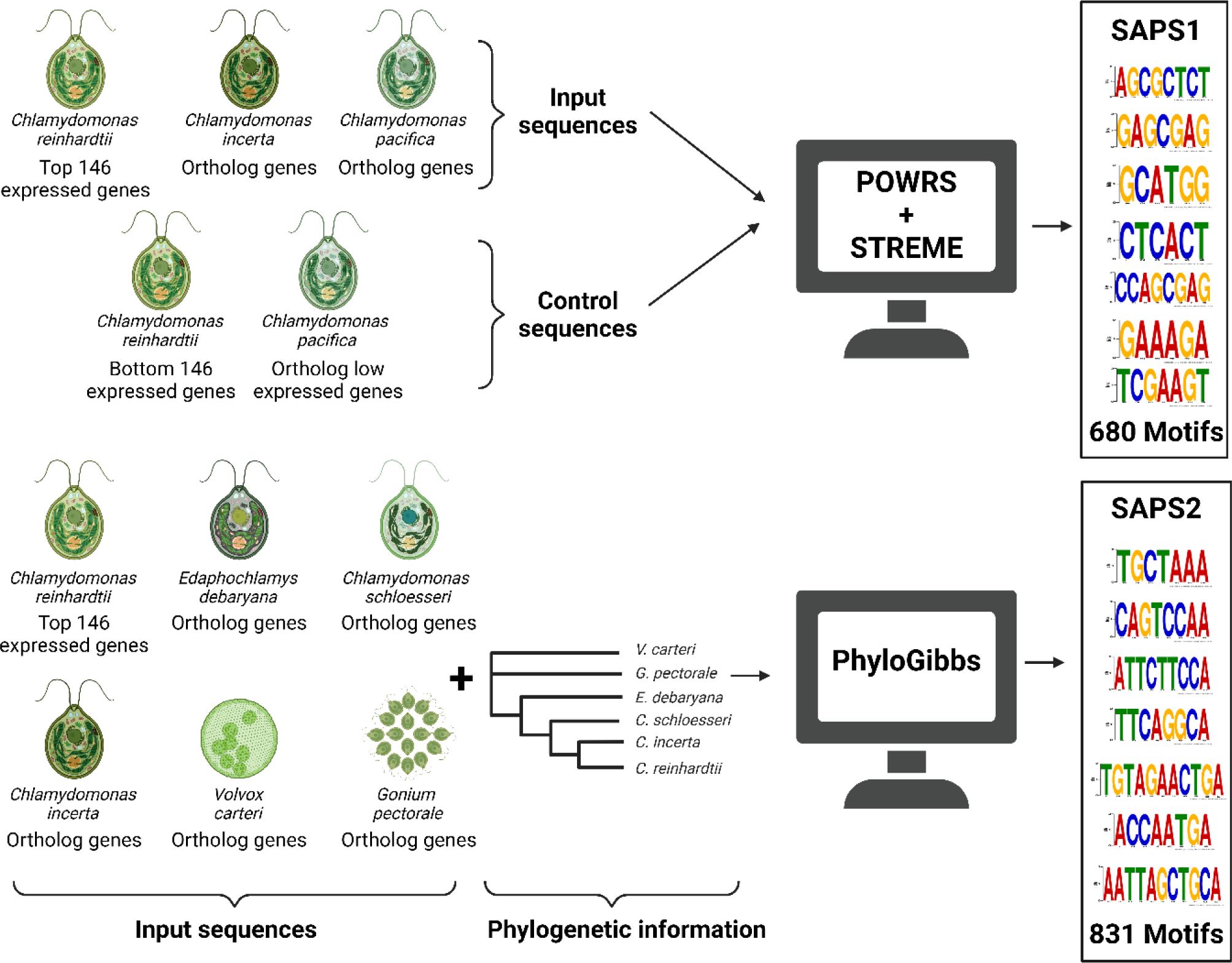
Identification and Utilization of CREs in SAPS1 and SAPS2 Libraries. Utilizing the top 146 highly expressed genes from *C. reinhardtii* and their orthologs in *C. incerta* and *C. pacifica*, alongside the least expressed 146 genes from *C. reinhardtii* and and their orthologs in *C. pacifica* as controls, 680 CREs were predicted via POWRS and STREME for SAPS1 promoters. For SAPS2, the same approach applied to *C. reinhardtii*’s highly expressed genes and orthologs in *E. debaryana, C. schloesseri, C. incerta, V. carteri,* and *G. pectoral*, incorporating phylogenetic distances, resulted in 831 CREs through PhyloGibbs.

For SAPS2, the same 146 promoter sequences from *C. reinhardtii* were used as input sequences, but the orthologue sequences used were those of *E. debaryana, C. schloesseri*, *C. incerta*, *V. carteri* and *G. pectoral*e. These species were selected to cover a spectrum of evolutionary distances, ranging from the closely related *C. incerta* and *C. reinhardtii*, to more distant species like *V. carteri*, with a divergence comparable to that between humans and chickens, estimated at 310 million years ago (Supporting Information – S1). PhyloGibbs predicts DNA motifs by combining over-representation analysis with evolutionary conservation across multiple sequence alignments of orthologous sequences. It uses a Bayesian model to discern functional binding sites, factoring in phylogenetic relationships, and applies simulated annealing alongside Monte-Carlo Markov chain sampling to rigorously evaluate and assign probabilities to these sites. This resulted in predicting 831 different CREs (Figure 1 and Supporting Information – S2).

### Insertion of putative cis-regulatory DNA elements into a synthetic promoter driving Zeocin resistance and GFP expression

The synthetic promoter chassis (SPC) was designed as a random 200 bp sequence with the following two constraints: the high GC content should resemble that of promoters in *C. reinhardtii* as described in Scranton et. al^10^, and the sequence should not contain any of the putative CREs to be tested in SAPS1 and SAPS2.

We engineered the synthetic promoters by inserting the predicted CREs in tandem arrays of 1, 2, or 3 copies. The insertion sites for these CREs were standardized across all promoters. Specifically, promoters with a single CRE had this element positioned at −104 relative to the transcriptional start site, with a variation for this position of ± 2bp. For promoters containing two CREs, the elements were positioned at −104 ± 2bp and −76 ± 2bp. Lastly, promoters with three CRE copies had their CREs located at −132 ± 4bp, - 104 ± 2bp, and −76 ± 2bp (Figure 2). This positioning was designed to keep approximately 20 bp of spacing between CREs to reduce steric hindrances between transcription factors binding to the promoter. This resulted in 2040 synthetic promoters in SAPS1 and 2493 synthetic promoters in SAPS2. These were subsequently cloned into a vector named pLibrary, culminating in the formation of two libraries: Synthetic Algal Promoters 1 (SAPS1) and 2 (SAPS2).

**Figure 2:**
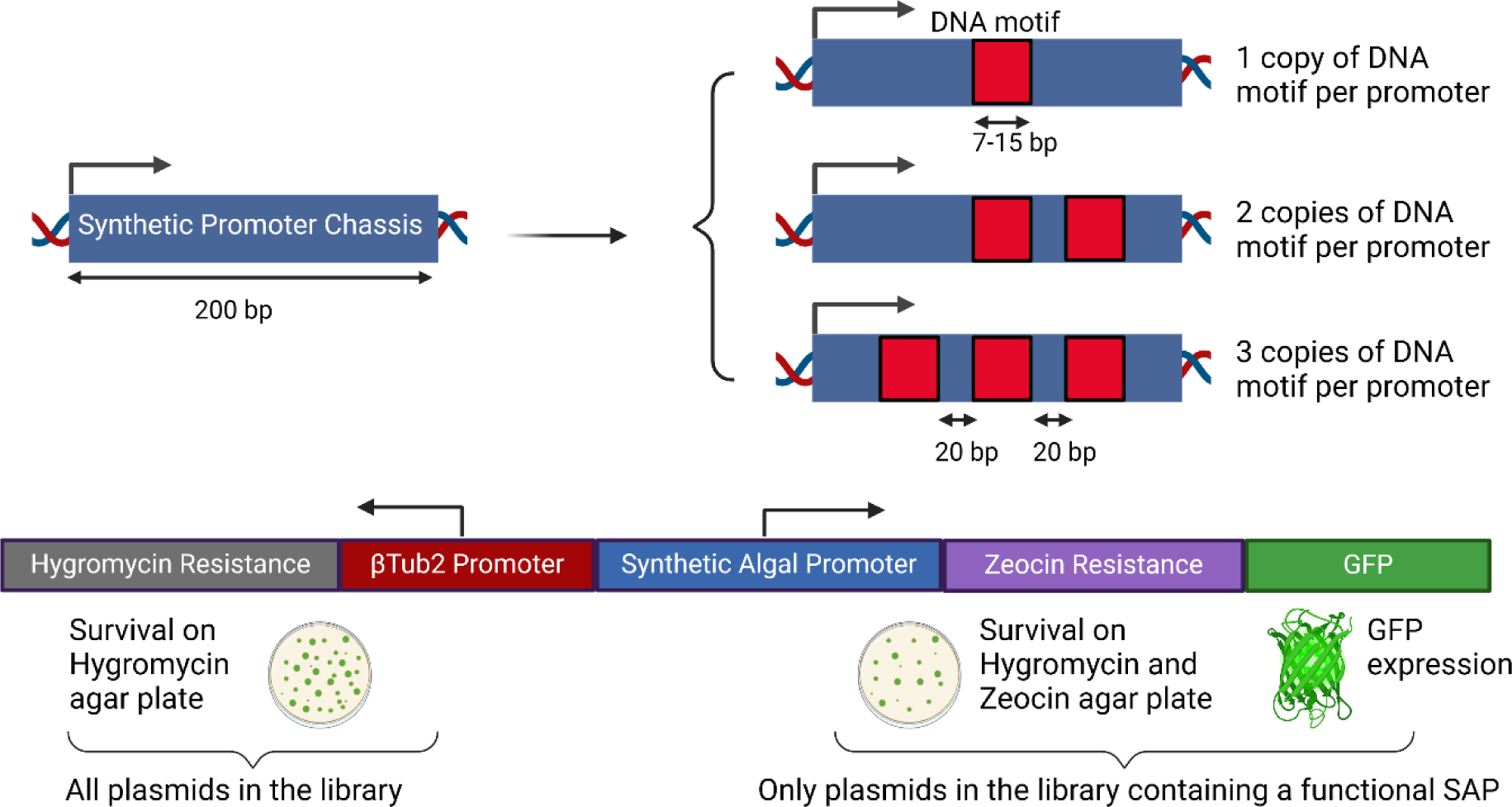
Design of Synthetic Promoter and Expression Vector. This figure illustrates the construction of synthetic promoters using a generated 200 bp DNA sequence with specific constraints. Predicted CREs were integrated in 1, 2, or 3 tandem copies at positions −104, −76, and −132 from the start site. The spacing between motifs was standardized to 20 bp. These synthetic promoters were positioned upstream of a GFP-fused zeocin resistance gene for expression analysis. For comparison, a β-Tubulin-2 promoter was utilized to regulate hygromycin resistance, serving as a control.

The synthetic promoters were made to drive the antibiotic resistance gene fused to the reporter gene GFP. This configuration allowed the selection of functional synthetic promoters through Zeocin resistance. Additionally, the GFP signal provides a measurable indication of the transcriptional activity of each promoter.

A high-throughput strategy for comparing the transcriptional activities of synthetic promoters within the same library was developed, combining differential antibiotic selections and NGS. This approach is predicated on the assumption that synthetic promoters demonstrating high transcriptional activity will sufficiently drive the expression of the ble gene to enable a significant proportion of transformants to surpass the zeocin survival threshold. Consequently, synthetic promoters with high transcriptional activity will have a higher relative abundance within the algal population post-transformation and zeocin selection. To accurately measure the relative abundance of each synthetic promoter within the algal libraries, we incorporated a secondary selection gene (APHVII) that imparts Hygromycin resistance. This gene was controlled by the well-established endogenous β-Tubulin 2 promoter^18^, conferring equal hygromycin-resistant capacity to all plasmids within both SAPS libraries. We strategically positioned this secondary selection gene upstream of the synthetic promoter, with transcription heading away from the synthetic promoter. This placement was essential to mitigate any promoter trapping effects that might influence the transcriptional activity of a synthetic promoter, especially if it were to integrate immediately downstream of a strong endogenous promoter during transformation of the synthetic promoter construct into the genome. Furthermore, to prevent transcriptional readthrough from the Hygromycin resistance cassette into the synthetic promoter, we encoded the cassette on the opposite DNA strand relative to the synthetic promoter (Figure 2).

Under hygromycin selection, each synthetic promoter‘s relative abundance should simply reflect its initial abundance within the physical DNA library utilized for cell transformation. The relative abundance under hygromycin and zeocin selection should provide insights into the transcriptional activity of the synthetic promoter driving the *ble* gene, as only strong synthetic promoters should allow sufficient transcription of the *ble* gene to enable ble resistance. Nonetheless, this measure is influenced by the inherent abundance of each promoter in the physical DNA library, as well. To account for this, the data is normalized by dividing the relative abundance under hygromycin and zeocin selection by the relative abundance under hygromycin alone, deriving what we termed the “HygZeo ratio”. This ratio indicates the extent to which a synthetic promoter is over- or underrepresented in the hygromycin and zeocin condition compared to hygromycin alone, which can be used as a relative measurement of the transcriptional activity of each synthetic promoter compared to the rest of promoters within the library.

### Assessment of synthetic promoter activity and CRE copy effect in *Chlamydomonas reinhardtii* using zeocin selection and next-generation sequencing

The SAPS1 and SAPS2 libraries were transformed into *C. reinhardtii* using electroporation. Post-transformation, each sample was divided into two parts: 10% of the volume was plated on TAP agar plates with 30 µg/mL Hygromycin (Hyg), and the remaining 90% on plates with 30 µg/mL Hygromycin and 10 µg/mL Zeocin (Hyg/Zeo). To ensure full representation of our promoter libraries in the high throughput in vivo testing, we established the goal of obtaining over 200,000 total transgenic lines (colonies) per library. This target assumes that only 1% of the obtained colonies are likely to exhibit significant transgene expression, due to factors such as random integration of transgenes into the nuclear genome leading to positional effects, RNA gene silencing, exonuclease activity causing truncated or damaged transgenes, and other epigenetic effects causing gene silencing^19^.

After performing over 60 transformations per library, around 280,000 SAPS1 colonies and 180,000 SAPS2 colonies were obtained on Hyg plates. The numbers on Hyg/Zeo plates were 12,000 colonies for SAPS1 and 14,000 colonies for SAPS2. Analyzing these figures and factoring in the 1:9 split in each transformation, it was calculated that only 0.47% of SAPS1 clones and 0.86% of SAPS2 colonies that survived in Hyg could also endure in Hyg/Zeo conditions. This means that even though only 12,000 and 14,000 colonies were obtained on the Hyg/Zeo plates, the actual number of colonies that were screened for transgene expression was much higher: 2,800,000 for SAPS1 and 1,800,000 for SAPS2. These numbers are in accordance with what has been previously reported in the literature^19^ for this type of selection. Additionally, this large number of transformants gives us confidence in our ability to thoroughly investigate these promoter libraries.

The algal colonies were separately pooled for each condition: SAPS1 in Hyg, SAPS1 in Hyg/Zeo, SAPS2 in Hyg, and SAPS2 in Hyg/Zeo, and these were then cultured in TAP liquid cultures. Subsequently, we took aliquots from each pool for genomic extraction and purification. We then amplified the synthetic promoter region of the integrated plasmid using PCR, as depicted in the Supporting Information – S9. The NGS process yielded approximately 150 and 120 million reads for SAPS1 and SAPS2, respectively. We processed and analyzed this data using the Galaxy platform^20^, obtaining a detailed examination of the promoter distribution within each sample.

Each processed read was matched to its corresponding promoter sequence. We then calculated the relative abundance of each promoter within SAPS1 and SAPS2 under the conditions Hyg and Hyg/Zeo, expressing the results as a percentage (Figure 3). Promoters show an average relative abundance of 0.05% in the initial Hyg library, aligning with expectations for 2,000 evenly distributed promoters (Figure 3 A). As previously explained, for each library, we divided the relative abundance of each promoter in the Hyg/Zeo condition (Figure 3 B) by their relative abundance in the Hyg condition, obtaining the normalized score “HygZeo ratio” (Figure 3 C). Using this metric, we ranked the promoters within the SAPS1 and SAPS2 libraries, from highest to lowest, being the number 1 promoter the one with the highest score and thus, the one with the highest transcriptional activity (Supporting Information – S2). In Figure 3, each promoter is depicted according to its relative abundance, with the most abundant ones positioned nearest to the graph’s origin. This arrangement, which places promoters in a descending order of abundance across the graph, effectively illustrates the distribution patterns of the entire promoter library tested. In the Hyg condition for both libraries (Figure 3 A), the distributions display a striking similarity between the mean and median values (SAPS1 mean: 0.049, median: 0.046, std: 0.026; SAPS2 mean: 0.040, median: 0.034, std: 0.028), indicative of a predominantly symmetrical distribution with a minor skewness towards higher values, likely attributable to a limited number of promoters exhibiting notably elevated relative abundances within the physical DNA library. Under the HygZeo condition (Figure 3 B), both libraries exhibit a marked departure from their previous symmetrical distribution, with a pronounced shift towards skewness (SAPS1 mean: 0.049, median: 0.004, std: 0.159; SAPS2 mean: 0.040, median: 0.006, std: 0.114). This is characterized by a drastic reduction in the relative abundance of most promoters, while a select few display a substantial increase in their relative abundance. When the relative abundances of each synthetic promoter in the HygZeo condition are normalized by dividing them by their corresponding abundances in the Hyg condition, we obtain the HygZeo ratio (Figure 3 C).

**Figure 3:**
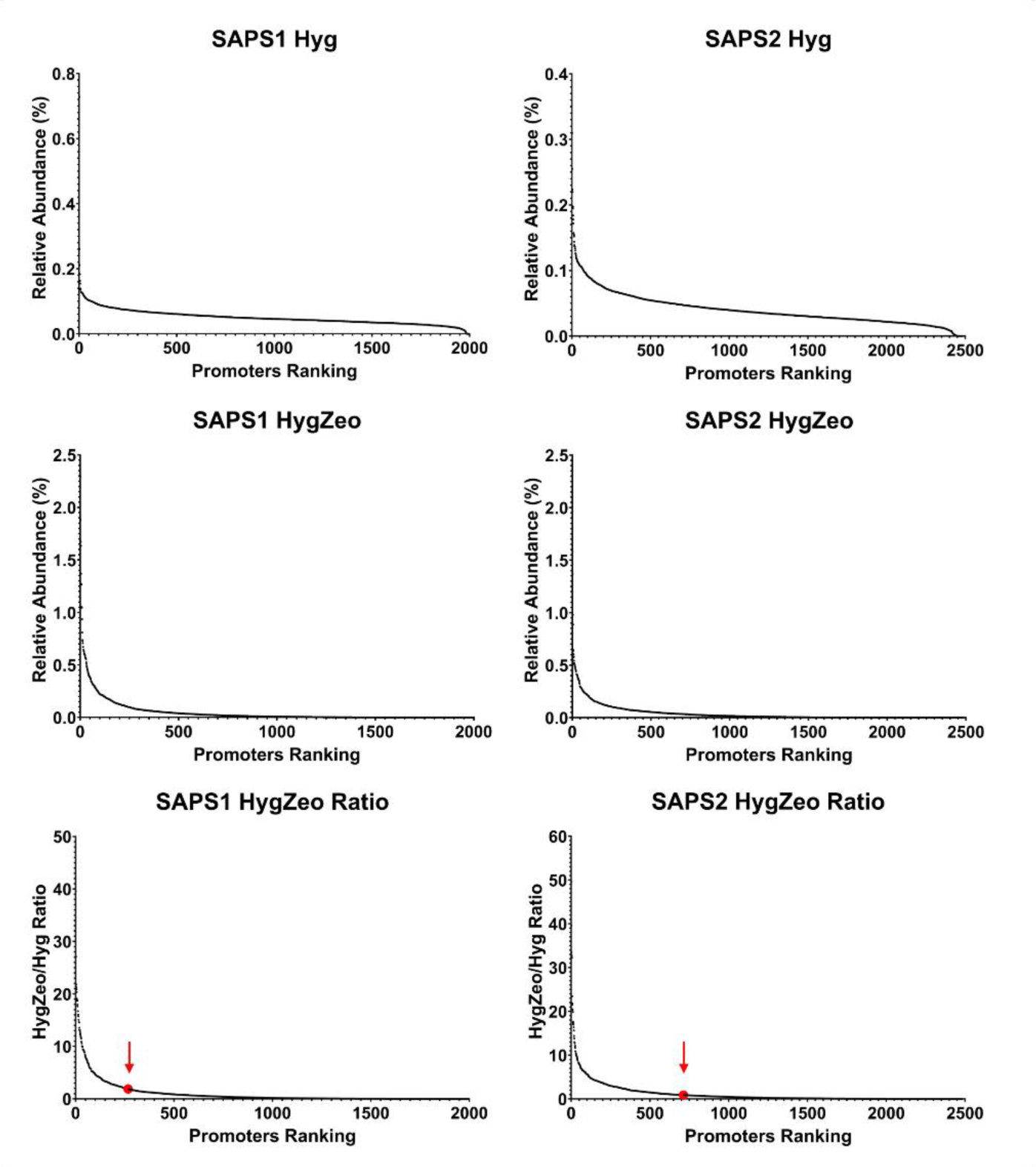
High Throughput Evaluation of the Synthetic Promoter Libraries Through Relative Abundance Measurements. The transformants resulting from the transformation of SAPS1 and SAPS2, selected on either hygromycin (Hyg) or hygromycin and zeocin (HygZeo), were pooled in liquid cultures. Through PCR, their promoter sequences were amplified and analyzed through NGS. The relative abundances, calculated as percentage of reads per promoter compared to the total amount of reads, are shown. Promoters are ranked based on their relative abundance, with the promoter showing the highest abundance in each graph receiving the ranking of 1. The relative abundance of each promoter under HygZeo selection is divided by the relative abundance of the same promoter under Hyg selection, obtaining the HygZeo ratio. The HygZeo ratio of the SPC is highlighted in red, with a red arrow indicating its position.

Analysis reveals that in both the SAPS1 and SAPS2 libraries, around 80% and 75% of the promoters, respectively, have a HygZeo ratio below 1 which indicates a reduction in their relative abundance when exposed to Zeocin. Moreover, the SPC without added CREs was also present in both libraries, and it ranked 266^th^ out of 2040 in SAPS1 and 712^th^ out of 2493 in SAPS2. This can be observed in Figure 3 C, in which the position of the SPC is indicated with a large red dot and a red arrow pointing at it. From this data, it can be assumed that the promoters with a significantly higher HygZeo ratio than that of the SPC contain a functional CRE. Promoters with the highest HygZeo ratio scores were selected that were at least three times the population’s standard deviation above the SPC HygZeo ratio. As a result, 37 promoters for SAPS1 and 40 for SAPS2 were identified, each exhibiting a relative abundance increase exceeding nine-fold, over the empty promoter construct (Supporting Information – S2). The best performing promoters present a relative abundance increase of 58-fold for SAPS1 and 49-fold for SAPS2, highlighting the dramatic effect that some CREs had on transcriptional activity, and hence Zeocin resistance, on the synthetic promoter that contained them.

The use of scoring of the promoters based on their relative abundance in HygZeo selection compared to Hyg alone, has advantages and disadvantages. The advantages are that this analysis allows for simple and fast high throughput screening of hundreds of thousands of individual transformation events, and thousands of functional promoters. The disadvantage is that it is susceptible to noise. The first source of noise is the fact that transgenes are randomly integrated into the nuclear genome of *C. reinhardtii*, which allows for the possibility of a synthetic promoter landing in a highly permissible place and overestimating its transcriptional strength, and vice versa. To reduce this noise, we cloned the Hyg resistance cassette upstream from the synthetic promoter and in the opposite DNA chain to avoid readthrough. Another possible source of noise is the fact that a percentage of transgenes integrated into the genome may integrate as truncated vectors. In this case, it would be possible for a promoter to have a damaged *ble* gene but still be counted in the Hyg sequencing sample which would underestimate its true increase in relative abundance in the HygZeo sample. Additionally, research suggests that a significant percentage of *C. reinhardtii* transformants may contain multiple insertions^21^. This would lead to inaccurate scoring as both promoters present in the transgenic cell would be measured, and thus assumed promoter activity could be a combination of their individual transcriptional activities.

We inserted CREs into synthetic promoters in 1, 2, or 3 tandem copies so that we could identify both basic transcriptional activity of a CRE, as well as explore any potential dose-dependent effect. This concept suggests that a greater number of CRE copies could lead to enhanced transcriptional activity, a hypothesis supported by previous research^22^. Analyzing the ranked synthetic promoters from each SAPS library, differentiated by the number of CRE copies, revealed a notable trend (Figure 4 A). Promoters containing a single CRE copy tended to cluster around the distribution median, indicating moderate transcriptional activity. In contrast, promoters with two CRE copies demonstrated a broader distribution, with transcriptional activity deviating from the median, both upwards and downwards. This divergence was even more pronounced in promoters with three CRE copies (Figure 4 A). When the top 25% of promoters from each library were isolated and separated by copy numbers, a significant difference in the means of their distributions was observed (Figure 4 B). As the copy number of CREs increased, a reduction in the mean of these populations was noted, indicating that promoters with a higher number of CRE copies generally exhibit enhanced transcriptional activity. Conversely, analysis of the bottom 25% ranked promoters revealed an increase in the mean of this population with rising copy numbers, suggesting that certain CREs may serve as repressor binding sites, where an increase in their copy number results in diminished promoter transcriptional activity (Figure 4 B).

**Figure 4:**
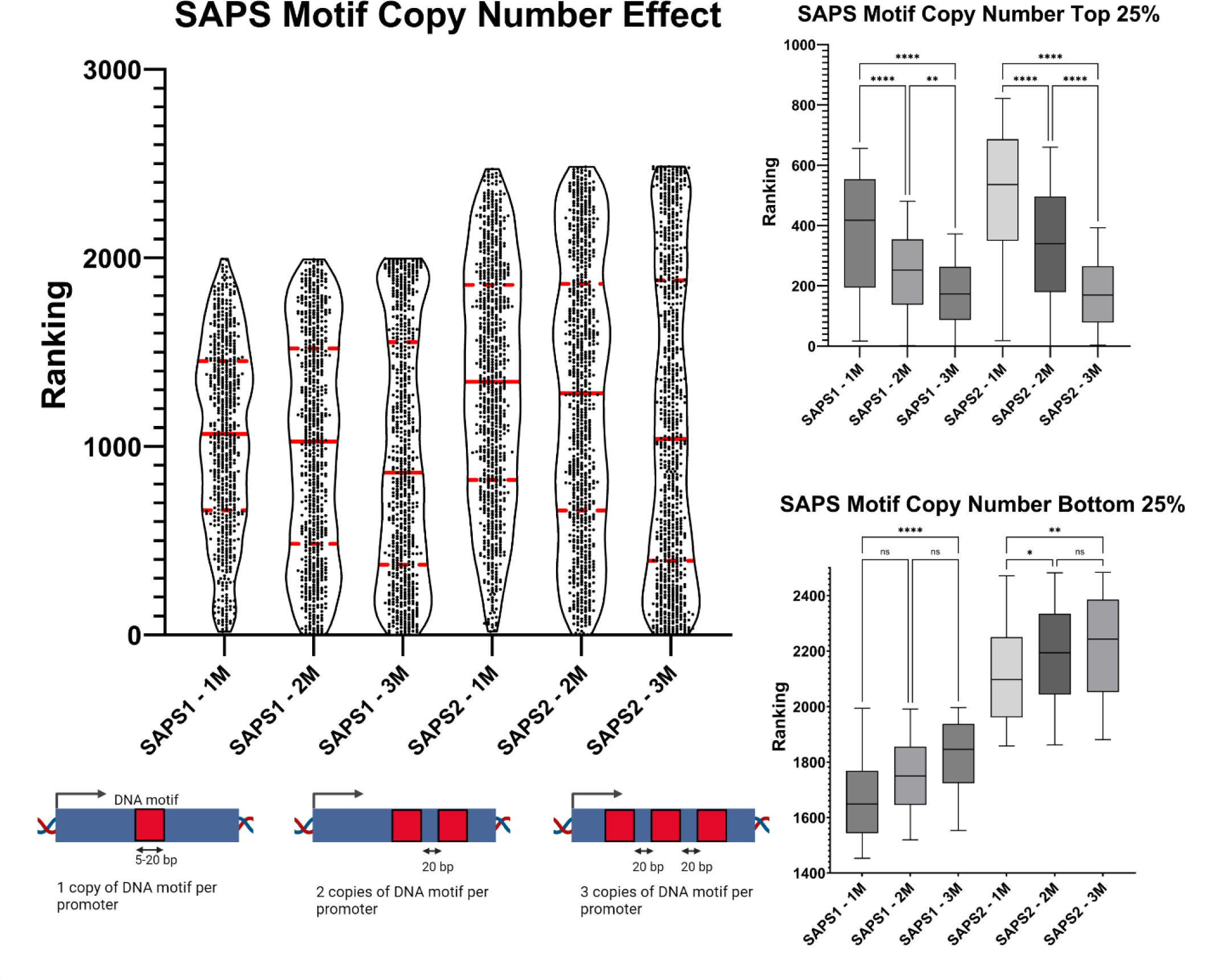
Assessing the Impact of Motif Copy Number on Synthetic Promoter Efficiency. This figure presents a truncated violin plot illustrating synthetic promoters’ performance, categorized by motif copy number and library, as measured by their HygZeo ratio rankings. The rankings are based on the HygZeo ratio, with rank 1 indicating the highest ratio. A continuous red line marks the median value, while dashed red lines denote the quartile divisions. Additional insets display the top 25% (upper quartile) and bottom 25% (lower quartile) of data points in a box-and-whiskers format, employing Tukey’s method for whisker calculation to identify outliers. The statistical significance of differences in median values across groups was determined using the Kruskal-Wallis test, followed by post-hoc analysis with Dunn’s multiple comparisons correction. Significance levels are indicated as follows: * for p < 0.05, ** for p < 0.01, *** for p < 0.001, and **** for p < 0.0001.

These findings suggest that our initial hypothesis is likely correct, in that increases in copy number of individual CREs either enhance (transcriptional activation) or diminish (transcriptional repression) the transcriptional activity of the synthetic promoter in which they are inserted. The proximity of the CRE to the transcription start site is recognized as critically important in native promoters^22^. It is conceivable that the optimal activity of the CRE is due to the positioning of the third motif copy at −132 base pairs, rather than the presence of additional CRE copies. The extensive range of CREs evaluated is anticipated to counterbalance the impact of this confounding variable. Although the positional effect likely influences some promoters, it is improbable that optimal performance across all CREs would occur at the −132 bp site. The insertion sites for all CREs were kept constant to limit experimental variables, despite the acknowledged significance of their positions. Accounting for variable positions would have increased the number of synthetic promoters to test to an unmanageable number.

### Selection of algal transformants containing synthetic promoters showing high GFP expression as measured via FACS

To validate the findings obtained from our high throughput promoter scoring system, we decided to evaluate the effectiveness of the synthetic promoters based on their ability to drive the expression of GFP. We identified several hundred transformants from each library harboring a synthetic promoter able to drive high levels of GFP. We then sequenced those individual transgenic lines to identify the sequence of the promoters, and compared those results to the ones obtained through the HygZeo ratio. The algal transformants that harbored either the SAPS1 or SAPS2 libraries, and were selected on Hyg or Hyg/Zeo, were subsequently combined in liquid cultures. From each pool of transformants, we isolated the top 5% of GFP expressors using FACS, and then plated those cells on Hyg or Hyg/Zeo plates. After the cells grew to form colonies, we picked 960 colonies, from each library and condition, into ten 96-well plates containing TAP media, and allowed them to grow for several days. Then, we measured their GFP signal in a plate reader, which we normalized by dividing it over their chlorophyll fluorescence as a measure of general cell growth.

In Figure 5 we show the normalized GFP signals compared to 96 clones from a strain producing GFP driven by the strong native AR1 promoter. From the figure we can observe that upon selection of SAPS solely on Hyg, significant GFP expression is observed in some cases, while most transformants exhibit minimal, if any, GFP expression. Transformants from SAPS2 library on Hyg in general exhibited much higher GFP expression than those from the SAP1 library. Upon selection of the libraries on Hyg/Zeo, both libraries showed a significant shift toward higher GFP expression, with certain transformants exhibiting GFP expression levels that surpassed those of the native AR1 promoter. This observation aligns with our expectations and corroborates the findings presented in Figure 3, which indicated that while most transformants harboring a SAPS promoter lack a highly functional promoter, a small subset possesses promoters with substantially enhanced transcriptional activity.

**Figure 5:**
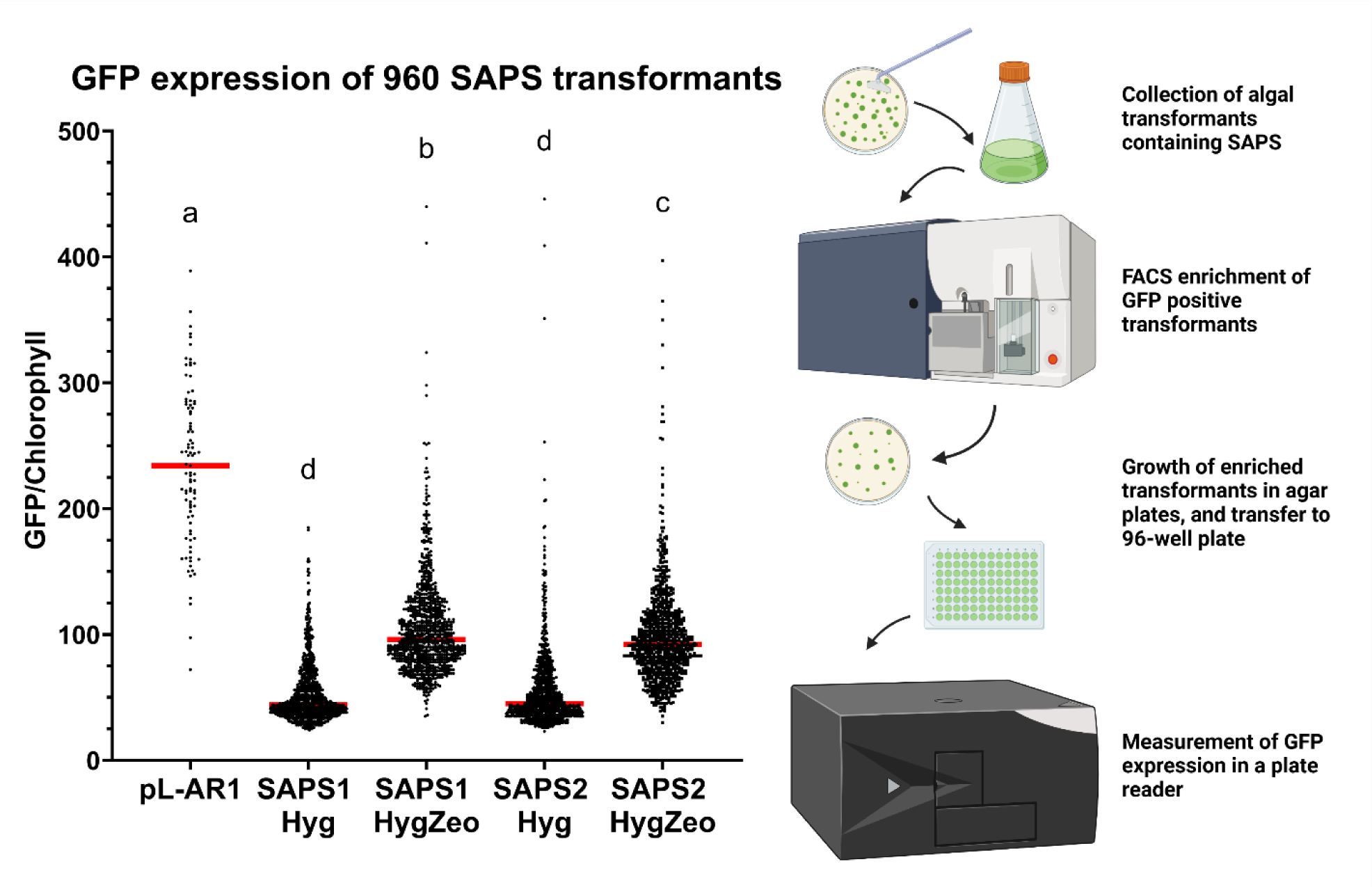
Analysis of the GFP expression of 960 SAPS transformants. The collected transformants for each library and condition were enriched for high GFP expression by sorting the top 5% GFP events. The cells were plated and inoculated into 96-well plates, allowed to grow for a couple of days and measured in a plate reader. GFP fluorescence, normalized against chlorophyll fluorescence, was measured in a plate reader (excitation: 505/6 nm, emission: 536/20 nm for GFP; excitation: 440/9 nm, emission: 680/20 nm for chlorophyll). Each sample has 960 data points, except for the positive control that contains 96 data points belonging to a clonal strain expressing GFP using the pL-AR1 vector. The statistical significance of differences in median values across groups was determined using the Kruskal-Wallis test, followed by post-hoc analysis with Dunn’s multiple comparisons correction. Samples annotated with different letters are statistically different from each other, with an adjusted p-value below 0.05.

960 individual transformants were measured on a plate reader for GFP expression, and several of the best GFP expressing transformants of SAPS1 and SAPS2 from HygZeo selection were individually sequenced to identify the synthetic promoter responsible for the high GFP expression. This resulted in a total of 115 SAPS1 and 254 SAPS2 transformants being sequenced, with promoters’ sequences being overrepresented in those with elevated HygZeo scores. (Supporting Information – S3). Out of those, 20 of the transformants sequenced for SAPS1 and 32 for SAPS2 belong to promoters with a HygZeo ratio 3 standard deviations above the mean. The frequency in which promoters with such HygZeo ratio appear in high GFP sequenced individuals is over 9-fold and over 8-fold higher than it would be expected from a random selection of promoters. These findings validate the HygZeo scoring shown in Figure 3 and provide us with a list of synthetic promoters containing predicted CREs shown to be able to drive the gene ble and GFP to high levels. In Table 1 we present a list of such synthetic promoters, described by the CRE they contain and the number of copies.

**Table 1.**
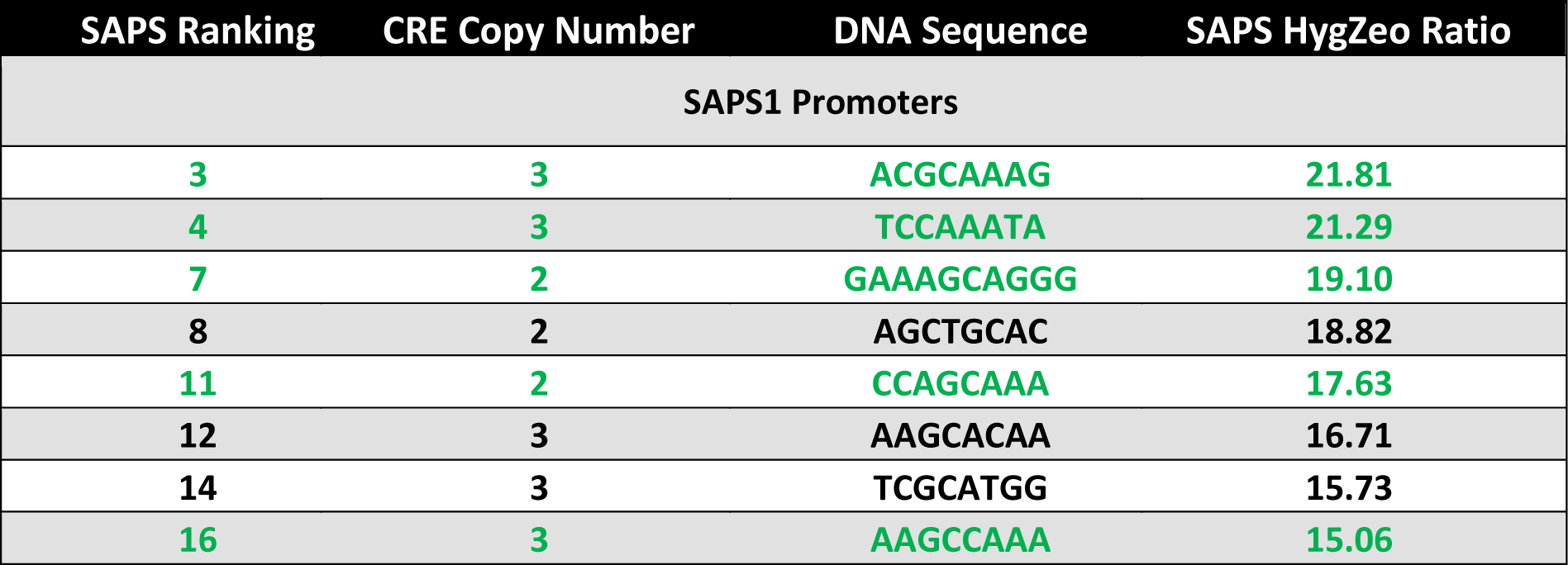
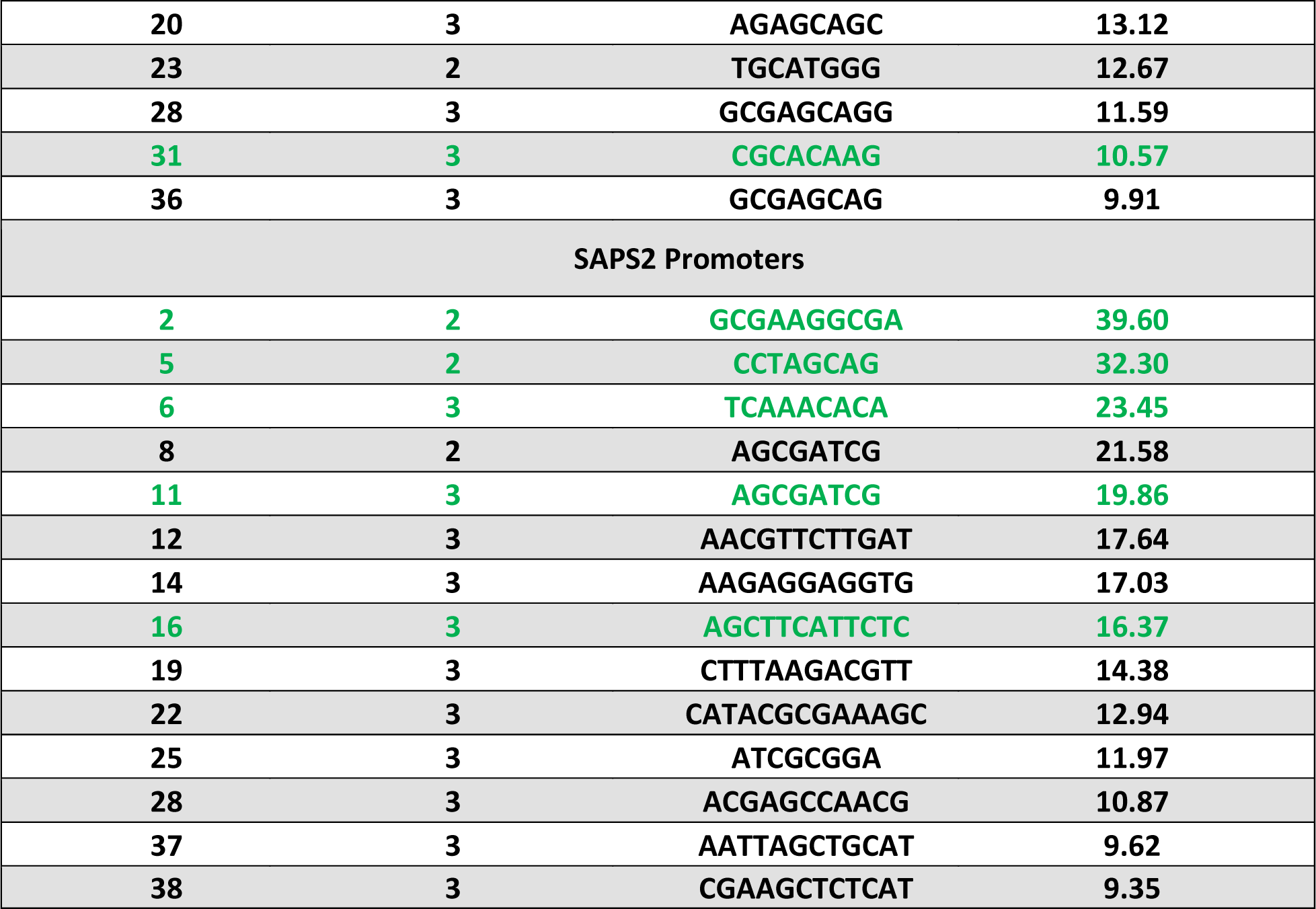
Validated Synthetic Promoters containing predicted CREs. This table shows synthetic promoters that showed a HygZeo ratio higher than three standard deviations above the SPC and were identified in individually sequenced high GFP expressors. Highlighted in green are promoters that were individually tested, with results shown in Figure 7.

| SAPS Ranking | CRE Copy Number | DNA Sequence | SAPS HygZeo Ratio |
| --- | --- | --- | --- |
| SAPS1 Promoters |  |  |  |
| 3 | 3 | ACGCAAAG | 21.81 |
| 4 | 3 | TCCAAATA | 21.29 |
| 7 | 2 | GAAAGCAGGG | 19.10 |
| 8 | 2 | AGCTGCAC | 18.82 |
| 11 | 2 | CCAGCAAA | 17.63 |
| 12 | 3 | AAGCACAA | 16.71 |
| 14 | 3 | TCGCATGG | 15.73 |
| 16 | 3 | AAGCCAAA | 15.06 |
| 20 | 3 | AGAGCAGC | 13.12 |
| 23 | 2 | TGCATGGG | 12.67 |
| 28 | 3 | GCGAGCAGG | 11.59 |
| 31 | 3 | CGCACAAG | 10.57 |
| 36 | 3 | GCGAGCAG | 9.91 |
| SAPS2 Promoters |  |  |  |
| 2 | 2 | GCGAAGGCGA | 39.60 |
| 5 | 2 | CCTAGCAG | 32.30 |
| 6 | 3 | TCAAACACA | 23.45 |
| 8 | 2 | AGCGATCG | 21.58 |
| 11 | 3 | AGCGATCG | 19.86 |
| 12 | 3 | AACGTTCTTGAT | 17.64 |
| 14 | 3 | AAGAGGAGGTG | 17.03 |
| 16 | 3 | AGCTTCATTCTC | 16.37 |
| 19 | 3 | CTTTAAGACGTT | 14.38 |
| 22 | 3 | CATACGCGAAAGC | 12.94 |
| 25 | 3 | ATCGCGGA | 11.97 |
| 28 | 3 | ACGAGCCAACG | 10.87 |
| 37 | 3 | AATTAGCTGCAT | 9.62 |
| 38 | 3 | CGAAGCTCTCAT | 9.35 |

We observed significant differences in the performance between the synthetic promoters from SAPS1 and SAPS2. At first, it appears that the synthetic promoters from SAPS2 were generally more effective than those from SAPS1. This can be observed in the percentage of colonies obtained under Hyg selection compared to those under HygZeo selection. For SAPS1, only 0.47% of the colonies survived in HygZeo compared to those that survived in Hyg alone, and that number elevates to 0.86% in the case of SAPS2. This difference in performance can also be seen in Figure 2, where 266 out of 2040 SAPS1 show a higher HygZeo ratio than the SPC compared to the 712 out of 2483 SAPS2. Moreover, in Figure 4 it can be observed that the effect exerted on the transcriptional activity of the promoters by the number of CREs inserted is steeper in SAPS2. All of these indicate that SAP2 library has more functional CREs than the SAP1 library. However, when we measured the GFP expression of the libraries, SAPS1 promoters appear to perform slightly better. This can be seen in Figure 5, where the overall GFP expression of the transformants tested for SAPS1 HygZeo was slightly superior to that of SAPS2 HygZeo. Thus, it seems the SAPS2 library has more functional CREs than the SAPS1 library, but that SAPS1 library has CREs that outperforms SAPS2 CREs when GFP expression is measured.

Selected transformants, one from SAPS1 and two from SAPS2, were subjected to imaging using a fluorescence microscope and compared with a GFP expressing clone containing the positive control promoter AR1. As shown in Figure 6, the cells exhibit red autofluorescence attributed to chlorophyll, while GFP, which is localized to the nucleus as a GFP:Ble fusion protein, can be visualized as a small green dot in the middle of the cells. The observed phenotypes in a selected subset of clones containing SAPS indicate that, while AR1 promotes GFP expression in a robust and uniform manner, these specific SAPS clones drive GFP expression at higher levels but with notable heterogeneity and induce a stressed phenotype, evidenced by cell enlargement in the affected cells. This stress may be linked to the significant accumulation of recombinant GFP within the nucleus, though such observations are not necessarily representative of all SAPS-containing transformants.

**Figure 6.**
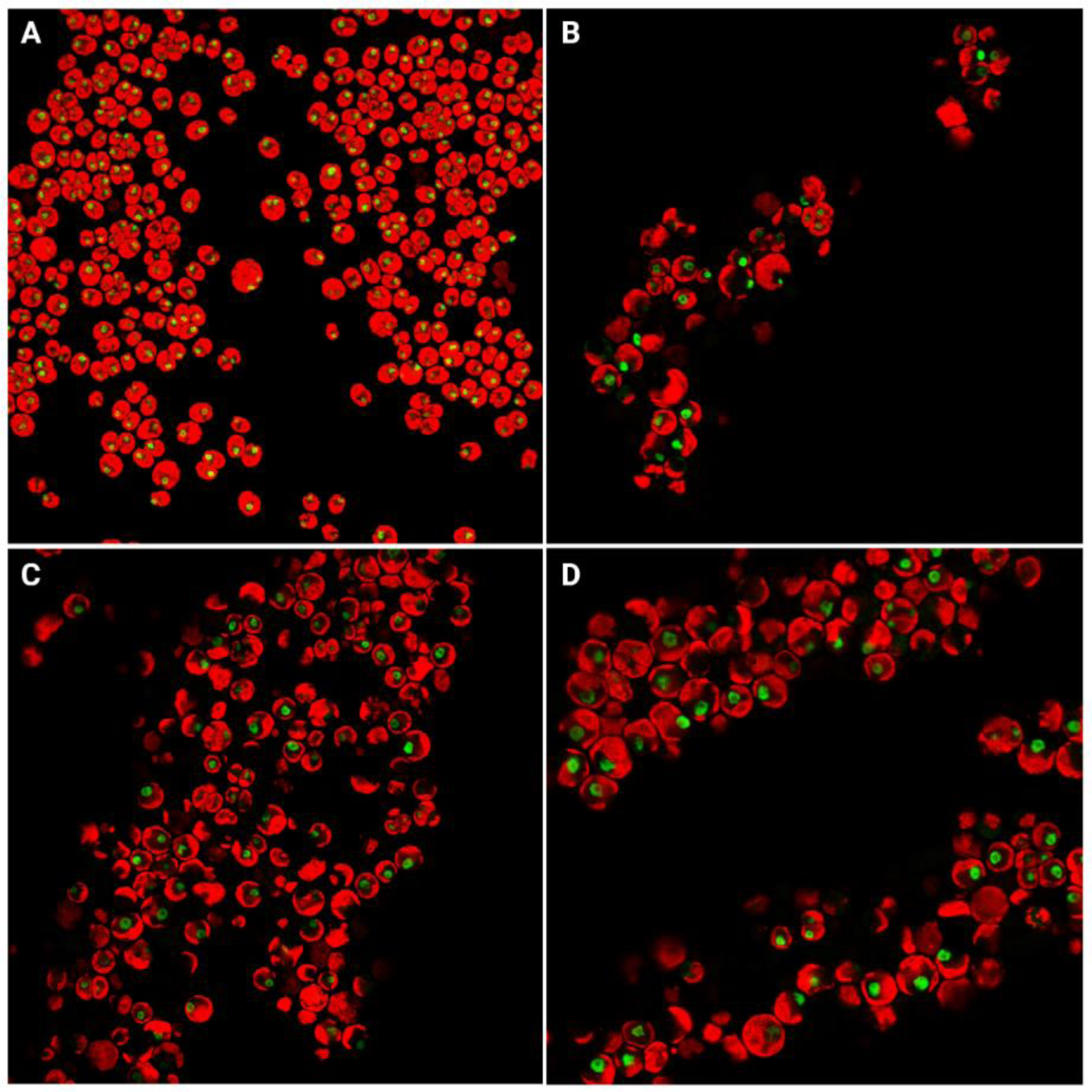
Differential GFP Expression in Synthetic Promoter Transformants via Fluorescence Microscopy. This image displays GFP levels in clonal cells, each with distinct promoters: A: pL-AR1, B: SAPS1 example, C-D: SAPS2 examples. Captured on a Leica DMi8 Inverted Confocal Microscope at 400X optical magnification with 0.75X digital zoom. Red indicates chlorophyll autofluorescence (excitation 555 nm, emission 617 nm); green signifies GFP (excitation 484 nm, emission 525 nm).

### Comparison of transcriptional activity of individual synthetic promoters as measured by GFP expression

To further validate our findings and accurately quantify the transcriptional activity of novel synthetic promoters, individual transformant with synthetic algal promoters were subjected to further analysis. A selection of fourteen synthetic promoters was made based on a combined assessment of those exhibiting a Hyg/Zeo ratio significantly above that of the SPC and the data from individually sequenced diverse synthetic promoters, highlighting a strategic integration of both datasets to identify promoters with varied anticipated transcriptional strengths. Those promoter constructs were individually transformed into *C. reinhardtii*, and their transcriptional activity was assessed by their GFP expression quantified through FACS of a population of transgenic lines from the single promoter. We also measured their transformation efficiency as a metric for transcriptional activity given that others have used this metric to evaluate the utility of a promoter in vivo^11^ and the fact that our HygZeo ratio can be thought of as a relative transformation efficiency score.

In Figure 7 A we can see the transformation efficiencies of all 14 SAPS compared to the two control promoters: a vector with the same design as those carrying synthetic promoters but containing only the SPC without motifs inserted (pL0), and an identically designed vector carrying the native AR1 promoter (pL-AR1). As shown, pL0 shows minimal transcriptional activity but is still able to generate colonies under Zeocin selection, with a transformation efficiency of approximately 57 colony forming units (CFUs) per µg of DNA. In contrast, the well-characterized and robust promoter AR1 showed a transformation efficiency of approximately 3000 CFUs/µg of DNA under the same conditions. Comparatively, the SAPS chosen for this experiment show transformation efficiencies ranging from approximately 500 to 3000 CFUs/µg of DNA. This data suggests that in terms of transformation efficiencies, some of the SAPS are as good as AR1, a strong chimeric endogenous promoter. As a direct comparison with pL0, we see that its transformation efficiency is improved by 53-fold with the addition of 3 tandem copies of the motif AGCTTCATTCTC. Other notable examples include the addition of 3 tandem copies of motifs TCCAAATA, ACGCAAAG and AGCGATCG, where the transformation efficiency was increased 41, 35 and 30-fold, respectively. All the SAPS tested yielded superior transformation efficiencies compared to pL0, except for pL1_70_1, which contained a single copy of the motif GGTACGGC.

**Figure 7.**
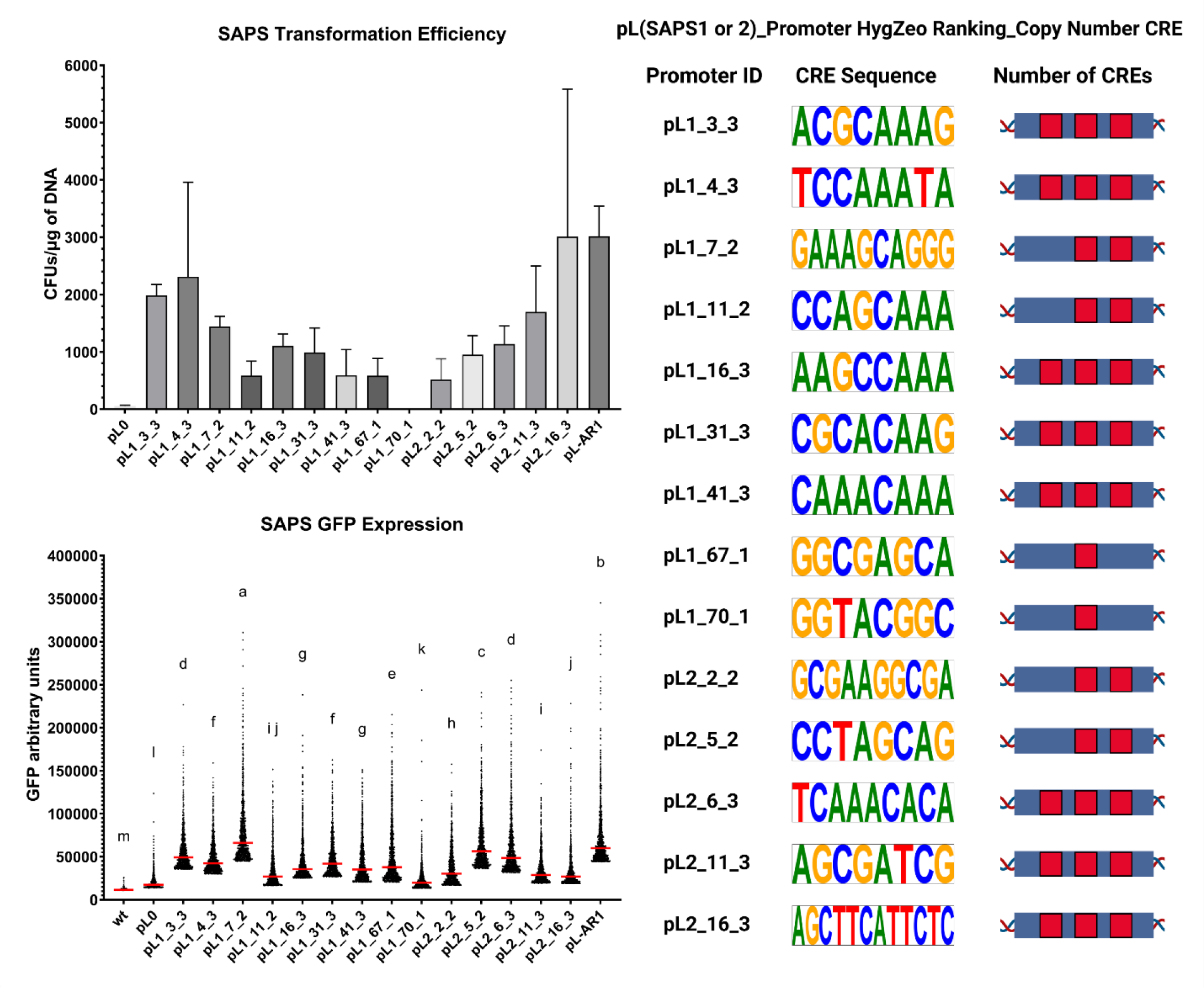
Evaluation of individual synthetic promoters. Top panel: Transformation efficiency of synthetic promoters, measured in CFUs using 1 µg of DNA for each, performed in triplicates, with pL0 and pL-AR1 as reference promoters. Bottom panel: GFP expression analyzed via FACS, pooling colonies from three transformations, showcasing the top 1000 GFP events per promoter, including wild-type (wt) cells for comparison. Statistical analysis was conducted with the Kruskal-Wallis test and Dunn’s post-hoc correction; promoters with differing letters signify statistically significant differences (adjusted p-value < 0.05). The promoter IDs structure is specified on the top right part of the figure: the first indicates the SAPS origin (1 or 2), the second number represents the HygZeo Ratio ranking within their library, and the third number specifies the CRE copy number for that promoter.

In Figure 7 B, the transcriptional activity of individual promoters, as measured by GFP expression in the top 1000 registered events during FACS analysis, is depicted. It is observed that pL0 exhibits minimal GFP activity, with a median GFP expression marginally above the background, as evidenced by comparison with the wild-type sample. All SAPS displayed a median GFP expression exceeding that of pL0, with improvements ranging from 1.4-fold to 9.2-fold (after background subtraction). pL-AR1 was observed to have a median GFP expression 8.2-fold higher than that of pL0. Among the SAPS evaluated, five exhibited GFP expression levels comparable to, or in one case surpassing, those of AR1. The SAPS, along with the CREs they contain, listed in descending order of GFP expression, are as follows: pL1_7_2 with 2 copies of GAAAGCAGGG, pL2_5_2 with 2 copies of CCTAGCAG, pL1_3_3 with 3 copies of ACGCAAAG, pL2_6_3 with 3 copies of TCAAACACA, and pL1_67_1 with 1 copy of GGCGAGCA. It was observed that the top two SAPS tested harbored an identical DNA sequence within their CREs, specifically the motif AGCAG, with both containing two copies. Additionally, pL1_67_1 possesses the motif AGCA that becomes an AGCAG due to the occurrence of a G immediately downstream from its insertion site. This notable occurrence suggests that the AGCAG motif may function as a transcription factor binding site.

## Conclusions

We now have a functional list of CREs that are available to build synthetic promoters for any green algae species. The natural next step is to combine different CREs in different combinations to explore potential synergistic effects and determine how far can transgene expression be pushed used these synthetic promoters. As an additional next step, we will test these CREs to identify those that increase gene expression under constant light and constant dark for *C. reinhardtii*, but this same procedure could be used to test any number of different growth conditions. In this way, CREs that respond to different environmental cues could be engineered into a single promoter for finely tune gene regulation in *C. reinhardtii*, and in any other algae species in which this strategy is deployed.

Advances in synthetic biology tools can be used to engineer cells to more efficiently produce a desired bioproduct, but this information can also be used to obtain a complete understanding of how the molecular machinery of the cell interacts with genetic information to yield different phenotypes. As the physicist Richard Feynman said, “What I cannot create, I do not understand”, we strive to grasp how genetic regulation works in microalgae by creating new genetic tools. In this work we strove to generate a comprehensive list of putative CREs that could then be used to create synthetic promoters of multiple functions, and we believe the list here is a very good start to building that tools necessary to take algae biotechnology to a new level.

## Methods

### Bioinformatic analysis and synthetic promoter chassis design

All genome and annotation data from NCBI ^23^. The data to determine the highest expressed genes can be found at the NCBI Gene Expression Omnibus repository under accession number GSE71469 (GSE71469_ChlamydomonasSynchronousDiurnalExpressionRPKM.txt.gz)^24^. To ensure the binding sites were still functional in short vectors and to include all core promoter elements, which are known to exist 200bp upstream of the transcriptional start site, The promoter length was approximated to be 500bp upstream of the translational start site. Furthermore, given its more reliable annotation, the translational start site was chosen instead of the Transcriptional Start Site as the reference location.

#### i. SAPS1 CRE prediction

Two algorithms were used to predict the CREs in SAPS1: POWRS and STREME^25^. As input sequences, the promoter sequences of the top 146 highest expressed genes in *C. reinhardtii* were selected, based on its diurnal transcriptome. The lowest 146 expressed genes were used as control sequences (Supporting Information – S1). We used Reciprocal Best Hit and BLASTN (e-value < 10^-5^) algorithms^26^ to identify orthologues of the selected genes in *C. incerta* and *C. pacifica* (Supporting Information – S1). The latest species is a novel extremophile green alga from the genus Chlamydomonas that was isolated at the University of California San Diego campus. In total, the number of orthologs for the 146 highly expressed genes from *C. reinhardtii* were: 146 genes in *C. pacifica* and 158 genes in *C. incerta*. For the lowest expressed genes, 116 ortholog genes were found in *C. pacifica*. The algorithm POWRS was run with parameters of ‘min_genes’ 120, ‘window_width’ 8, ‘cluster_limit’ 800 and ‘seed_size’ 8. STREME was executed using the parameters ‘minw’ 6 and ‘maxw’ 15, and only motifs that were 8 base pairs or more identical between at least two of the following species: *C. reinhardtii*, *C. incerta*, or *C. pacifica*, were chosen. In both cases, only the motifs that were found in at least two of the three species and showed p-value smaller than 10^-2^ were selected. This resulted in the prediction of 680 CREs (Supporting Information – S2).

#### ii. SAPS2 CRE prediction

The algorithm PhyloGibbs^27^ was used to predict the CREs in SAPS2. The top 146 highest expressed genes in *C. reinhardtii* were used as input sequences, as well as the orthologue genes for the highest 146 from C. reinhardtii found in the species *Edaphoclamys debaryana* (90 genes), *Chlamydomonas schloesseri* (132 genes), *C. incerta* (158 genes), *Volvox carteri* (4 genes) and *Gonium pectorale* (92 genes) (Supporting Information – S1). DiAlign^28^ was utilized, following the parameter guidelines recommended by Siddhartan et al.^27^, for deducing the multiple sequence alignments of the promoter regions across the top 146 genes in *C. reinhardtii* and their orthologs. The phylogenetic relationships among these species were elucidated using Benchmarking Universal Single-Copy Orthologs (BUSCO) genes^29^. Multiple sequence alignments were conducted with MUSCLE^30^, gene trees were inferred via CLUSTALW^31^, and the species tree was constructed using ASTRAL III^32^. This methodology resulted in the identification of 831 CREs.

#### iii. Synthetic Promoter Chassis

The synthetic promoter chassis (SPC), a 200bp sequence on which CREs are embedded, was designed in-silico using a random sequence generator based on probabilities derived from Scranton et al^10^. The first 60 bp have probabilities for ACGT of [0.2, 0.25, 0.35, 0.2], while the next 140bp have probabilities of [0.28, 0.24, 0.2, 0.28]. After the sequence is generated, if any of the predicted CREs in SAPS1 and SAPS2 is present, the SPC sequence is regenerated to exclude the CRE. This process ensures SPC closely resembles the native core promoter of *C. reinhardtii* while avoiding interference from CREs during testing.

### Expression Vector design

The expression vector is designed to have two open reading frames in opposite directions. One ORF contains a Hygromycin resistance cassette composed of: the *C. reinhardtii* β2-Tubulin promoter from −1 to −199 (both positions included), the 113 bp of the 5’ UTR of the same gene, the APH7 gene which confers *C. reinhardtii* with hygromycin resistance, and 455 bp of the 3’ UTR of the β2-Tubulin gene. The other ORF contains a zeocin resistance gene fused to the fluorescent protein GFP mclover. A promoter being tested, described below, is placed upstream of the 23 bp 5’ UTR of the *C. reinhardtii rbcs2* gene followed by the coding region, and ending with the 234 bp of the 3’ UTR of the *C. reinhardtii rbcs2* gene. The coding region, from start to stop codon, contains 1563 bp and is composed of several parts. There are two introns, rbcs2 intron 1 (positions 169 .. 313) and 2 (positions 927 .. 1255). The first protein is the CDS of the *ble* gene, codon optimized for the nuclear genome of *C. reinhardtii*, which confers resistance against the antibiotic zeocin. The second protein is the GFP variant mclover, which was also codon optimized. Finally, there is the 8 DYKDDDDK amino acid tag at the C-terminus.

The promoters used to drive the *ble*-GFP fusion were the AR1 promoter^33^ in the case of pLibrary-AR1 (pL-AR1), or the SPC previously described in the case of pLibrary-0 (pL0). Any tested SAPS were used as promoters replacing the SPC. Sequences fully annotated for the vectors pL0 and pL-AR1 can be found in the Supporting Information S5 and S6, respectively. The sequence of all the SAPS1 and SAPS2 assembled are also included (S7 and S8). The strain and annotated plasmid sequence for pL0, were deposited in the public instance of the Agile BioFoundry Inventory of Composable Elements (ICE)^34, 35^ Registry (https://public-registry.agilebiofoundry.org/folders/40) and are physically available upon reasonable request.

### Promoter Library synthesis and assembly into the vector

#### i. Promoter library synthesis and assembly into vector

Promoter libraries SAPS1 and SAPS2 were synthesized by Twist Bioscience (South San Francisco, CA), containing 2040 and 2493 variants, respectively. The sequences of all the SAPS designed can be found in the Supporting Information – S7 and S8. In brief, assembly of the promoter libraries into the pL0 vector was achieved by using Gibson Assembly®, reactions with PCR-amplified promoter libraries and pL0 backbone containing Gibson overhangs designed by j5^35, 36^ (j5.jbei.org). Following assembly, each library was transformed into *E. coli* via electroporation and subsequently extracted for downstream steps. Details of these steps are described below. Unless otherwise specified, all primers were synthesized by Integrative DNA Technologies (Coralville, IA), and all PCR reactions were set up with Q5 Master Mix (New England Biolabs, Ipswich, MA) according to the manufacturer’s protocol.

Promoter libraries SAPS1 and SAPS2 were amplified with j5-designed primers ‘barcode_pool_F’ (5’ TGCTGGAAGTGTCATAGCGCAAGAAAGNNNNNNNNGTCCTTCCCGGGCCAGGC 3’) and ‘barcode_pool_R’ (5’ TCTCTTGTAAAAAAGTAGTTGAGGATCCCCCACTTATTGCG 3’), and the following thermal cycler conditions: 98°C for 3 minutes, 10 cycles of 98°C for 20 seconds, 63°C for 30 seconds, 72°C for 40 seconds, and a final extension of 72°C for 3 minutes. SAPS1 and SAPS2 were amplified with 10 PCR cycles to limit amplification bias. The pL0 backbone was amplified using j5-designed primers ‘backbone_F’ (5’ TGGGGGATCCTCAACTACTTTTTTACAAGAGAAGTCACTCAACATC 3’) and ‘backbone_R’ (5’ TTTCTTGCGCTATGACACTTCCAGC 3’), using thermal cycler conditions consisting of 98°C for 3 minutes, 25 cycles of 98°C for 15 seconds, 63.5°C for 30 seconds, 72°C for 3.5 minutes, and a final extension of 5 minutes at 72°C. Following amplification, the backbone PCR reaction was digested overnight with DpnI (New England Biolabs, Ipswich, MA) at 37°C. All amplicons were then verified by gel electrophoresis and purified using a Wizard Gel Purification kit (Promega, Madison, WI).

Gibson assemblies of each promoter library were executed using NEBuilder Hifi Assembly Master Mix (New England Biolabs, Ipswich, MA) following the manufacturer’s protocol. Assemblies were then transformed into MegaX DH10β T1R Electrocomp™ Cells (Thermo Fisher Scientific, Waltham, MA), in 1 mm sterile cuvettes (VWR, Radnor, PA) with a Gene Pulser Xcell (Bio-Rad Laboratories Inc., Hercules, CA) at 2.0 kV, 200 Ω, and 25 µF. For each promoter library, four electroporation reactions were executed in parallel to increase the diversity of promoters transformed. The four reactions were pooled following recovery, and plated on a Bioassay Qtrays (Molecular Devices, San Jose, CA) containing 250 mL of 100 µg/mL Carbenicillin LB agar. After 18 hours of growth at 37°C, each promoter library’s Qtray was scrapped into 35 mL of 100 µg/mL Carbenicillin LB media using cell scrapers (Sarstedt, Newton, NC), and extracted using a Midiprep kit (Qiagen, Hilden, Germany).

#### ii. Library preparation and QC

To verify the assemblies and assess the distribution of promoter variants, promoter regions of each promoter plasmid library were amplified and ligated to sequencing adapters, then purified and sequenced using Next Generation Sequencing (NGS). This process is described below. Promoter regions of each plasmid library were amplified with primers ‘screen_F’ (5’ GGAAGTGTCATAGCGCAAGAA 3’) and ‘screen_R’ (5’ AGATGTTGAGTGACTTCTCTTGTAA 3’), and thermal cycler conditions consisting of 98°C for 30 seconds, 7 cycles of 98°C for 10 seconds, 54.8°C for 30 seconds, 72°C for 40 seconds, and a final extension of 72°C for 1 minute. The promoter region was amplified with 7 PCR cycles to limit amplification bias. Following verification with gel electrophoresis, amplicons were purified using a Wizard Gel Purification kit. Amplicons were then A-Tailed and ligated to xGen-UDI-UMI Adapters (Integrative DNA Technologies, Coralville, IA) using a KAPA HyperPrep PCR-Free kit (Roche, Basel Switzerland) according to the manufacturer’s protocol. The resulting libraries were purified twice at 0.9X using Ampure XP Beads (Beckman Coulter, Indianapolis IN), and quantified using a Qubit 3.0 Fluorometer (Thermo Fisher Scientific, Waltham, MA) and Qubit dsDNA High Sensitivity kit (Thermo Fisher Scientific, Waltham, MA). One nanogram of each library was then run on a Bioanalyzer 2100 (Agilent, Santa Clara, CA) with a High Sensitivity DNA Kit (Agilent, Santa Clara, CA) to assess quality and adapter ligation.

Each library was then sequenced on a Miseq (Illumina, San Diego, CA) using a MiSeq Reagent Kit v2 (500 cycles) with paired-end sequencing. Each library’s R1 and R2 reads were then interleaved into one FASTQ using BBTools reformat^37^. Lastly, sequencing data was analyzed by synbioqc-libqc^38^ to verify promoter pool assembly and quantify promoter variant diversity per promoter pool. Histograms visualizing the distribution of promoter variant counts per library are shown in Supporting Information – S10.

### Algal strain used and culture conditions

The algal strain used to perform this study was *Chlamydomonas reinhardtii* cc-1690. The algae was cultured in TAP media^39^, grown under continuous light with a photosynthetically active radiation of 125 μE/m2/sec, on a shaker table rotating at 125 RPM, and a constant temperature of 25 °C.

### Algal transformation and selection

The plasmid DNA was subjected to double digestion using KpnI and XbaI enzymes (New England Biolabs, Ipswich, MA, USA), followed by purification with the Wizard SV Gel and PCR Clean-up System (Promega Corporation, Madison, WI, USA), without fragment separation. DNA concentration was quantified using the Qubit dsDNA High Sensitivity Kit (Thermo Fisher Scientific, Waltham, MA, USA). The DNA amount for each transformation was adjusted to ensure 1 µg of payload DNA, based on the payload DNA to backbone DNA ratio. Transformation is performed through electroporation as described in this study^40^.

Transformant selection occurred on two types of plates. For library transformations, resuspensions of 1 mL in TAP were divided: 100 µL onto TAP agar with 30 µg/mL Hygromycin (Hyg plates) and 900 µL onto TAP agar with 30 µg/mL Hygromycin plus 15 µg/mL Zeocin (HygZeo plates). Individual promoter transformations involved plating the entire 1 mL resuspension on HygZeo plates.

### Next-Generation-Sequencing of the algal libraries

#### i. Genome Extraction

Genomic DNA was extracted from algal cells using SDS DNA extraction buffer (1% SDS, 20 mM Tris-HCl pH 8.0, 2 mM Sodium EDTA pH 8.0, 200 mM NaCl). Approximately 50 µL of packed cells from late-log phase culture were lysed in 1625 µL of this buffer at 65°C. The lysate was then subjected to extraction with an equal volume of Phenol:Chloroform:Isoamyl Alcohol pH 8.0 and centrifuged at 5000 rcf for 4 minutes. The aqueous phase was washed twice with 1875 µL of ethanol-stabilized Chloroform and once with an equal volume of Chloroform. DNA was precipitated from the final aqueous phase by adding an equal volume (750 µL) of isopropanol, followed by centrifugation at 16,000 rcf for 30 minutes at 4°C. The DNA pellet was washed thrice with 70% ethanol, air-dried, and resuspended in 150 µL of 1X TE pH 8.0. Quantification was performed using a Qubit HS DNA kit, yielding concentrations suitable for use in Q5 PCR reactions.

#### ii. Library Amplification

The libraries were prepared for sequencing by PCR using the extracted genomic DNA as template. The PCR protocol involved two amplification steps using New England Biolabs (NEB, Ipswich, MA, USA) reagents. In the first PCR, 500 ng of DNA genomic DNA was amplified over 13 cycles using a master mix containing Q5 Phusion Buffer, GC Enhancer, primers, dNTPs, and Q5 Polymerase from NEB, followed by gel purification. The primers used for this round were forward primer

TCGTCGGCAGCGTCAGATGTGTATAAGAGACAGgctggaagtgtcatagcgcaag and reverse primer GTCTCGTGGGCTCGGAGATGTGTATAAGAGACAGcccacttattgcgagactgggc.

The second PCR, with 20 cycles, utilized 5 ng of the purified amplicon from the first round of PCR per reaction with a similar master mix composition. This round included the addition of barcodes for posterior sequencing deconvolution. The primers used were:

SAPS1 Hyg Forward: AATGATACGGCGACCACCGAGATCTACACTCGTGGAGCGTCGTCGGCAGCGTC

SAPS1 Hyg Reverse: CAAGCAGAAGACGGCATACGAGATCGCTCAGTTCGTCTCGTGGGCTCGG

SAPS1 HygZeo Forward: AATGATACGGCGACCACCGAGATCTACACTATAGTAGCTTCGTCGGCAGCGTC

SAPS1 HygZeo Reverse: CAAGCAGAAGACGGCATACGAGATATATGAGACGGTCTCGTGGGCTCGG

SAPS2 Hyg Forward: AATGATACGGCGACCACCGAGATCTACACCTACAAGATATCGTCGGCAGCGTC

SAPS2 Hyg Reverse: CAAGCAGAAGACGGCATACGAGATTATCTGACCTGTCTCGTGGGCTCGG

SAPS2 HygZeo Forward: AATGATACGGCGACCACCGAGATCTACACTGCCTGGTGGTCGTCGGCAGCGTC

SAPS2 HygZeo Reverse: CAAGCAGAAGACGGCATACGAGATCTTATGGAATGTCTCGTGGGCTCGG

Post-amplification, the products were cleaned up, diluted to a uniform concentration of 80 ng/µL, and 25 µL of each were pooled, ensuring equal molar ratios for subsequent steps.

#### iii. Next-Generation Sequencing

For sequencing analysis, we utilized the services of Azenta (Chelmsford, Massachusetts, USA). Our sample was composed of equimolar parts of our four libraries as previously described. The libraries were designed to be compatible with Illumina’s sequencing platform, including standard Illumina adapter sequences, primer binding sites for both Read 1 and Read 2, as well as dual index sequences with a length of 10 base pairs each for sample deconvolution. We employed a sequencing configuration of 2×150 bp with 150 million reads, which fully covered our 222 bp amplicon and allowed 78 bp of overlapping sequences in the middle, and the read depth was more than enough to guarantee full coverage of our library.

#### iv. Sequence Data Processing

For NGS data analysis we employed a sophisticated Galaxy^20^ workflow for processing and analyzing sequencing data. The workflow began with the processing of paired-end raw reads, followed by adapter trimming and quality control measures. Next, the reads were merged based on overlap criteria. This step was crucial for reconstructing the complete sequences of our amplicons. Subsequently, specific genomic regions, namely promoters, were extracted from these merged reads. The extracted sequences were then aligned to a promoter database, facilitating the identification of promoter regions in our sample. Post-alignment, the data was converted into a suitable format for further analysis. The final steps of the workflow focused on extracting and sorting relevant read and sequence identifiers, culminating in a count of distinct reads and barcodes. This workflow enabled a comprehensive sequencing data analysis, ensuring accurate identification and quantification of genomic elements of interest. The workflow can be visited at the following link: https://usegalaxy.eu/u/yasintorres/w/synthetic-algal-promoter-library---count-analysis

#### v. CRE Abundance Analysis

Each synthetic promoter was correlated with its corresponding read count. The total reads for each library were summed to determine the library’s total reads. Relative abundance of each promoter (SAPS1 Hyg, SAPS1 HygZeo, SAPS2 Hyg, and SAPS2 HygZeo) in the library was calculated by dividing the individual promoter reads by the total library reads. Finally, to obtain the HygZeo ratio, the relative abundance of each promoter in the HygZeo condition was divided by its abundance in the Hyg condition, for both SAPS1 and SAPS2. Spreadsheets with the calculations can be found in the Supporting Information – S2. The results obtained were graphically represented using GraphPad Prism (GraphPad Software, San Diego, California, USA).

### In vivo CRE activity based on Fluorescence Activated Cell Sorting

In this study, we utilized Fluorescence-Activated Cell Sorting to analyze GFP expression in transformed *Chlamydomonas reinhardtii* cells in a similar manner as described in Sproles et. al^19^. The analysis was performed using a Sony MA900 Cell Sorter with 405 nm, 488 nm, 638 nm, and 561 nm lasers. Our gating strategy focused on cell size and chlorophyll autofluorescence, excitation laser 488 nm, and emission filter 695/50 nm to isolate single cells. We sorted and analyzed the top 5% of cells with the highest GFP expression, excitation laser 488 nm, and emission filter 525/50 nm. Data for 500,000 events per sample were collected in FCS files and then analyzed using FlowJo (FlowJo LLC, Ashland, Oregon, USA). The top 1000 GFP data points per sample were extracted, and graphs were generated using GraphPad Prism (GraphPad Software, San Diego, California, USA).

The cells were collected on 15 mL conical tubes containing 1 mL of TAP media. After sorting, the cells were plated onto HygZeo plates, and colonies were visible 3-5 days after.

### Fluorescent Confocal Microscopy

High-resolution confocal imaging was performed at the UCSD Microscopy Core using a Leica DMi8 Inverted Confocal Microscope (Leica Microsystems, Wetzlar, Germany). Samples were illuminated with a tunable White Light Laser, with excitation and emission wavelengths selected for chlorophyll (excitation: 555 nm, emission: 617 nm) and GFP (excitation: 484 nm, emission: 525 nm). The oil immersion 40x objective lens with a numerical aperture 1.3 and a digital zoom 0.75x was utilized. Laser power, gain, and pixel dwell time were adjusted to optimize image clarity and minimize photobleaching. Images were acquired with a resolution of 2048×2048 pixels, using Leica LAS X software (Leica Microsystems, Wetzlar, Germany). Spectral deconvolution was conducted within the software to mitigate fluorophore signal overlap, and channels were merged to assess fluorophore colocalization, ensuring precise and artifact-free data for subsequent analysis.

### Sequencing of individual transformants

Individual transformants grown on HygZeo plates were patched onto similarly treated plates with drawn grids for clone tracking. Subsequently, these transformants were picked into 96-well plates, each well containing 200 μL of TAP media supplemented with 10 μg/mL Zeocin. After a growth period of 3 days, measurements were conducted using a Tecan Infinite m200 Pro plate reader (Tecan Group Ltd., Männedorf, Zurich, Switzerland). Two key parameters were assessed: GFP fluorescence, indicative of promoter transcriptional activity, and chlorophyll concentration for cell number normalization. GFP was measured at an excitation wavelength of 505/9 nm and emission wavelength of 536/20 nm, with the plate reader set to a gain of 245. Chlorophyll measurements were taken at an excitation wavelength of 440/9 nm and emission wavelength of 680/20 nm, with a gain of 160. Transformants exhibiting the highest GFP-to-chlorophyll ratio were selected for further analysis.

For DNA extraction, the chosen transformants from the original HygZeo plates were resuspended in 20 μL of 10X Tris-EDTA (TE) Buffer and subjected to a boiling step at 95°C for 10 minutes. Following this, 2 μL of the resultant lysate was used as a template for PCR amplification. The PCR employed the primers: forward GATCACAAGCTCGAGTGGCC and reverse GACCGCGCTGATGAACAGGG. The amplified PCR products were then purified using a standard clean-up kit. Finally, these purified samples were forwarded to Eton Bioscience Inc. (San Diego, California) for Sanger sequencing analysis.

## Abbreviations

SAPS: Synthetic Algal Promoters
SPC: Synthetic Promoter Chassis
CRE: Cis-regulatory Elements
FACS: Fluorescence Activated Cell Sorting
NGS: Next-generation Sequencing
GFP: Green Fluorescent Protein
Hyg: Hygromycin
Zeo: Zeocin
CFU: Colony Forming Unit
pL0: pLibrary-0
pL-AR1: pLibrary-AR1
WT: Wild-type
POWRS: POsition-sensitive WoRd Set
STREME: Simple, Thorough, Rapid, Enriched Motif Elicitation

## Supporting Information

S1. DNA Input Sequences
Zip file containing the input dna sequences used for all three algorithms: POWRS, STREME and PhyloGibbs

S2. SAPS Data.xlsx
SAPS1 Hyg – Contains data regarding the counts for each SAPS1 under Hyg
SAPS2 Hyg – Contains data regarding the counts for each SAPS2 under Hyg
SAPS1 HygZeo – Contains data regarding the counts for each SAPS1 under HygZeo
SAPS2 HygZeo – Contains data regarding the counts for each SAPS2 under HygZeo
SAPS1 – Contains the compiled data for all SAPS1 counts and the calculated HygZeo ratios
SAPS1 – Contains the compiled data for all SAPS1 counts and the calculated HygZeo ratios

S3. Individually sequenced transformants.xlsx
All sequenced promoters – Contains the corresponding promoter ID for each individually sequenced transformant
Unique sequenced promoters - Contains the corresponding promoter ID for each unique sequenced promoter

S4. Plate reader data for SAPS transformants.xlsx
Each sheet contains the raw data for GFP expression of all the algal transformants analyzed

S5. pLibrary-0.gbk
GenBank DNA file containing the sequence of the plasmid pLibrary-0

S6. pLibrary-AR1.gbk
GenBank DNA file containing the sequence of the plasmid pLibrary-AR1.

S7. SAPS1 fasta sequences.txt Sequences of all SAPS1 in fasta format

S8. SAPS2 fasta sequences.txt Sequences of all SAPS2 in fasta format

S9. NGS Reads Annotated.docx
Docx file containing an image showing the structure of the NGS reads.

S10. Distribution of promoter variants in SAPS1 and SAPS2.docx
Docx file containing histograms showing the promoter distribution of each promoter library.

## Acknowledgements

The authors thank Ernst Oberortner, Lisa Simirenko, and Duncan Scott for writing and maintaining the synbioqc-libqc analysis repository. The authors also thank Jan-Fang Cheng, Ian Blaby, Robert Evans, and Garima Goyal for providing the promoter pool cloning and quality control workflow protocol that was adapted for this study. Additionally, the authors thank Rita Kuo for her insight and guidance in troubleshooting the promoter library prep and quality control process. The author(s) declare financial support was received for the research, authorship, and/or publication of this article by the United States Department of Energy through the grant DE-EE0009671 (APEX). We acknowledge The Scripps Research Institute Flow Cytometry Core Facility for the services provided, as well as the UCSD School of Medicine Microscopy Core which was funded through the grants P30 NS047101 and S10 OD030505. This work was part of the Agile BioFoundry (agilebiofoundry.org) supported by the U.S. Department of Energy, Energy Efficiency and Renewable Energy, Bioenergy Technologies Office through contract DE-AC02-05CH11231 between Lawrence Berkeley National Laboratory and the U.S. Department of Energy. The views and opinions of the authors expressed herein do not necessarily state or reflect those of the U.S. Government or any agency thereof. Neither the U.S. Government, nor any agency thereof, nor any of their employees makes any warranty, expressed or implied, or assumes any legal liability or responsibility for the accuracy, completeness, or usefulness of any information, apparatus, product, or process disclosed or represents that its use would not infringe privately owned rights. The U.S. Government retains and the publisher, by accepting the article for publication, acknowledges that the U.S. Government retains a nonexclusive, paid-up, irrevocable, worldwide license to publish or reproduce the published form of this manuscript, or allow others to do so, for U.S. Government purposes. The Department of Energy will provide public access to these results of federally sponsored research in accordance with the DOE Public Access Plan (http://energy.gov/downloads/doe-public-access-plan). Funding for open access charge: U.S. Department of Energy.

