## Supporting Information - S9 for "Bioinformatic prediction and high throughput in vivo screening to identify cis-regulatory elements for the development of algal synthetic promoters"

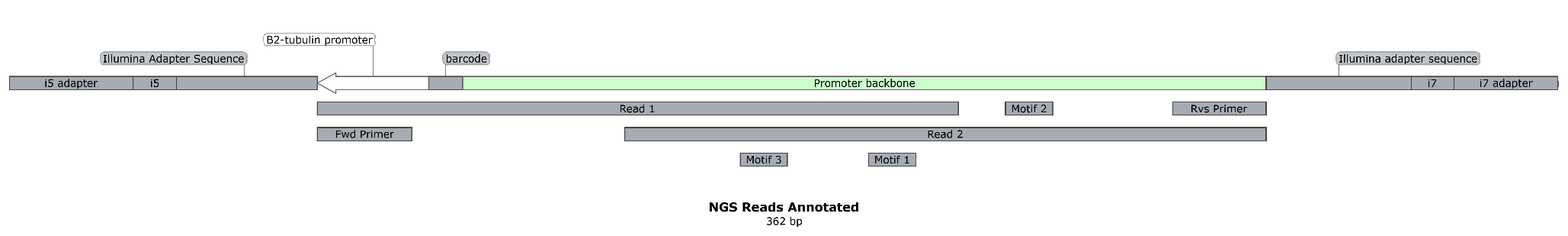

**S10. NGS Reads Annotated**. The promoter sequence was amplified through two rounds of PCR. The first round included the annealing primers Fwd and Rvs, with adapter overhangs. The second PCR introduced the i5 and i7 indexes and adapter sequences. The amplicons were sequenced in a 2x150bp Illumina Sequencing reaction, with 78 bp of overlap between paired-end reads.
