## Supporting Information - S10 for "Bioinformatic prediction and high throughput in vivo screening to identify cis-regulatory elements for the development of algal synthetic promoters"

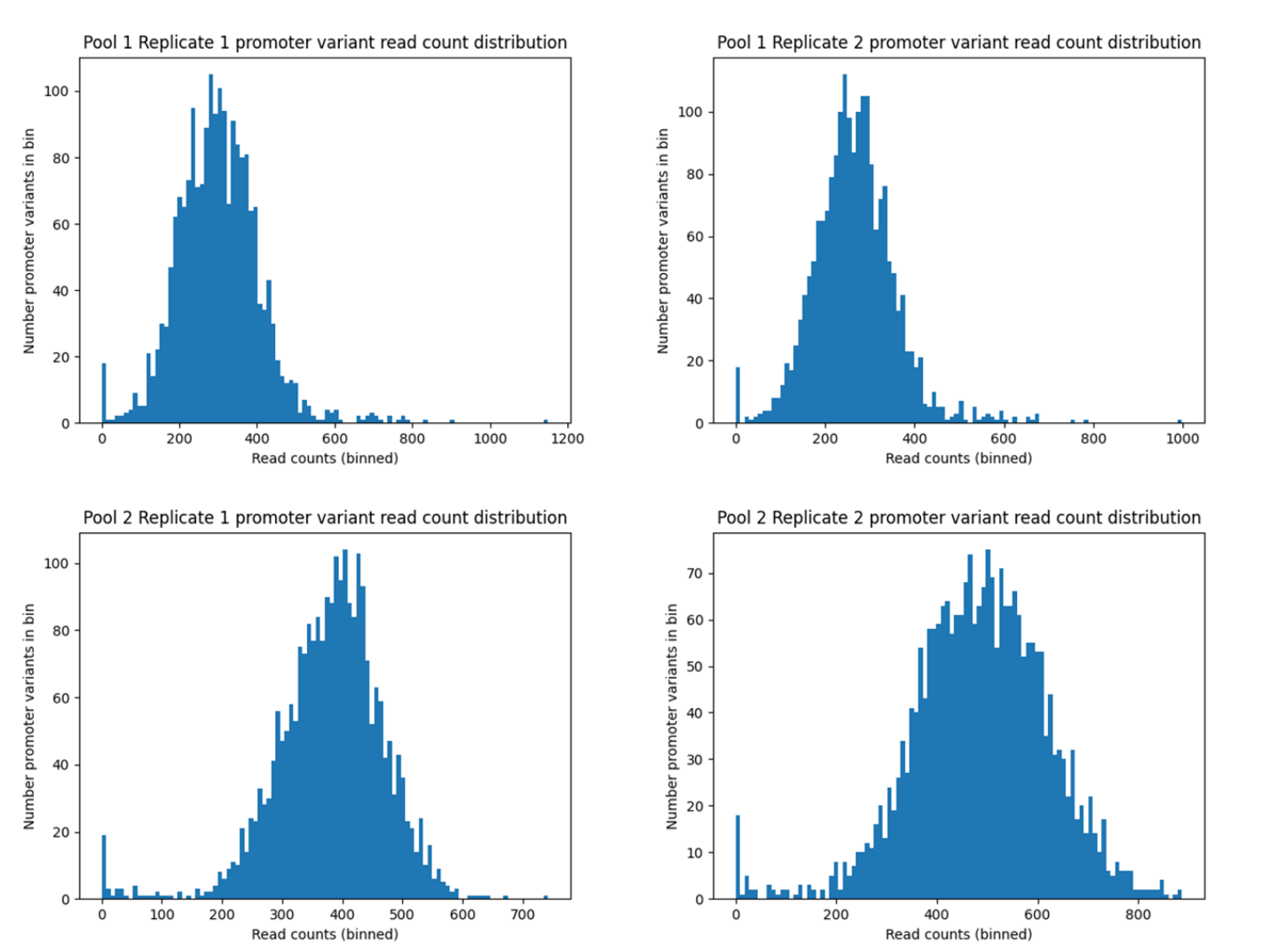

**S11. Distribution of promoter variants in SAPS1 and SAPS2**. Replicate batches of plasmid preparations of SAPS1 (pool 1) and SAPS2 (pool 2) were analyzed by sinbioqc-libqc to verify promoter pool assembly and quantify diversity of promoters within the libraries. Histograms visualizing the distribution of promoter variant counts per library as shown.
